# Oncogenic YAP mediates changes in chromatin accessibility and activity that drive cell cycle gene expression and cell migration

**DOI:** 10.1101/2022.09.08.507127

**Authors:** Maria Camila Fetiva, Franziska Liss, Dörthe Gertzmann, Julius Thomas, Benedikt Gantert, Magdalena Vogl, Nataliia Sira, Grit Weinstock, Susanne Kneitz, Carsten P. Ade, Stefan Gaubatz

**Affiliations:** Theodor Boveri Institute and Comprehensive Cancer Center Mainfranken, Biocenter University of Wuerzburg, Wuerzburg, 97074, Germany

## Abstract

YAP, the key protein effector of the Hippo pathway, is a transcriptional co-activator that controls the expression of cell cycle genes, promotes cell growth and proliferation and regulates organ size. YAP modulates gene transcription by binding to distal enhancers, but the mechanisms of gene regulation by YAP-bound enhancers remain poorly understood. Here we show that constitute active YAP5SA leads to widespread changes in chromatin accessibility in untransformed MCF10A cells. Newly accessible regions include YAP-bound enhancers that mediate activation of cycle genes regulated by the Myb-MuvB (MMB) complex. By CRISPR-interference we identify a role for YAP-bound enhancers in phosphorylation of Pol II at Ser5 at MMB-regulated promoters, extending previously published studies that suggested YAP primarily regulates the pause-release step and transcriptional elongation. YAP5SA also leads to less accessible “closed” chromatin regions, which are not directly YAP-bound but which contain binding motifs for the p53 family of transcription factors. Diminished accessibility at these regions is, at least in part, a consequence of reduced expression and chromatin-binding of the p53 family member ΔNp63 resulting in downregulation of ΔNp63-target genes and promoting YAP-mediated cell migration. In summary, our studies uncover changes in chromatin accessibility and activity that contribute to the oncogenic activities of YAP.

## INTRODUCTION

The evolutionary conserved Hippo pathway plays important roles in development, cell proliferation, organ size control and cancer (1–4). The hippo cascade involves multiple kinases, such as MST1/2 and LATS which ultimately phosphorylate YAP and its close paralog TAZ, triggering their cytoplasmic retention and degradation via the proteasome (5). In contrast, when Hippo is inactive, unphosphorylated YAP/TAZ enter the nucleus and act as transcriptional coactivators by binding to TEAD transcription factors (1, 2, 6). Deregulation of the upstream hippo tumor suppressors can cause uncontrolled growth and cancer (3, 4). Aberrant activation of YAP is known to contribute to cancer initiation, progression and drug resistance and is generally correlated with a poor outcome (7).

Although YAP induces the expression of a number of target genes through binding to promoters, recent studies have shown that YAP primary regulates gene expression by binding to distal transcriptional enhancers, regulatory DNA elements that activate the expression of their distant target genes by the formation of chromatin loops (8–11). Enhancers are highly abundant in the mammalian genome and they play important roles in spatiotemporal regulation of gene expression (12). In cancer cells, enhancers have been shown to be critical to reprogram gene expression and to promote oncogenic activities (13–15). Deregulated enhancers also play a role in the establishment and maintenance of transcriptional addiction, a dependency on transcription factors and chromatin regulators for sustained proliferation of cancer cells (16). YAP is recruited to enhancers mainly by TEAD transcription factors, but is also known to cooperate with other transcription factors and co-activators. For example, many YAP-regulated enhancers contain both TEAD and AP-1 motifs where YAP synergizes with JUN/FOS to promote tumor cell proliferation and transformation (9, 17, 18). In addition, YAP interacts with the mediator and with the bromodomain protein BRD4 to promote assembly of the pre-initiation complex and trigger transcriptional elongation (19, 20).

In previous work we have shown that YAP cooperates with the B-MYB transcription factor to activate G2/M cell cycle genes (21, 22). B-MYB (also called MYBL2) binds to the MuvB core complex to form the Myb-MuvB (MMB) complex, which activates late cell cycle genes (23–27). Mechanistically, by binding to distant enhancers YAP promotes the association of B-MYB with MuvB, leading to the formation of the MMB-complex at the TSS of cell cycle genes and resulting in induction of target gene expression (21). In addition, YAP also enhances the expression of B-MYB, contributing to an increased rate of mitosis and hyperproliferation (21, 22, 28).

Here we investigated epigenetic changes by oncogenic YAP using untransformed human breast epithelial MCF10A cells expressing constitutive active YAP5SA. We found that YAP5SA leads to global chromatin changes resulting in thousands of newly opened and closed genome regions. By CRISPR-interference and ChIP, we identified a role for YAP in increasing levels of Ser5-phosphorylated RNA Pol II at the CDC20 promoter. ChIP and biochemical experiments demonstrate that YAP5SA leads to the enrichment of Ser5-phosphorylated RNA Pol II at the promoters of MMB-target genes and provide evidence that this modification is mediated by CDK7. We also demonstrate that YAP leads to closing and inactivation of enhancers bound by the ΔNp63 tumor suppressor. We show that the loss of ΔNp63 chromatin binding and downregulation of ΔNp63 target genes is critical for cell migration by oncogenic YAP.

## MATERIALS AND METHODS

### Cell lines

MCF10A-YAP5SA cells have been described previously (29). MCF10A cells were cultured in DMEM/F-12 supplemented with 5% horse serum, 1% penicillin/ streptomycin, 10 μg/ml insulin, 500 ng/ml hydrocortisone, 20 ng/ml EGF and 100 ng/ml cholera toxin. The expression of YAP5SA was induced in MCF10A-YAP5SA cells by the addition of 0.5 μg/ml doxycycline. MCF10A-YAP5SA-ΔNp63 and MCF10A-ER-YAP2SA-ΔNp63 cells were generated by infection with pINDUCER20-ΔNp63 lentivirus and selection with 1 mg/ml neomycin. For simultaneous induction of YAP5SA and ΔNp63, MCF10A-YAP5SA-ΔNp63 cells were treated with 0.2 μg/ml doxycycline.

### ATAC-Seq

100,000 MCF10A cells were washed with ice cold PBS and lysed in ATAC lysis buffer (10mM Tris pH 7.4, 10mM NaCl, 3 mM MgCl_2_, 0.1% Tween 20 and freshly added protease inhibitor cocktail (Sigma)) by incubating on ice for 10 minutes. Nuclei were collected by spinning at 500 g for 10 minutes at 4°C. The transposition reaction mix (25 μl 2X TD buffer, 2.5 μl TDE1 Nextera transposase (Illumina), 16.5 μl PBS, 0.5 μl 1% digitonin, 0.5 μl 10% Tween-20, and 5 μl of nuclease free water) was added to nuclei and incubated at 37°C for 30 mins. Next, the reaction was cleaned using the MinElute PCR purification kit (Qiagen), and a PCR with 10-13 cycles was performed using the NEBNext High Fidelity 2X PCR Master Mix (NEB) and Ad1_noMX and Ad2.1–2.12 barcoded primers described in (30). Size selection of the libraries was performed with Agencourt AMPure XP beads (Beckman Coulter). Library quality and fragment size distribution was analyzed on a fragment analyzer (Advanced Analytical). Paired end 2 × 75 bp sequencing was performed on the NextSeq 500 platform (Illumina).

### CUT&RUN

CUT&RUN was carried out as described (31, 32). Briefly, 500,000 cells were washed twice with wash buffer (20 mM HEPES pH 7.5, 150 mM NaCl and 0.5 mM spermidine containing protease inhibitor) and captured with 20 μl conacavallin A magnetic beads (Polyscience). Cells were resuspended with antibody buffer (wash buffer with 0.005% digitonin and 2 mM EDTA) and incubated with 2μg of YAP-antibody (Novus Biologicals, NB110-58358) for 2 hours at room temperature. Samples were then washed twice with wash buffer containing 0.005% digitonin and incubated with 700 ng/ml of purified protein-A/G-MNase fusion (pA/G-MNase) on a shaker at 4°C for 1 h followed by two more washes in digitonin wash buffer and once in low salt rinse buffer [20mM HEPES pH7.5, 0.5mM Spermidine, 0.005% Digitonin]. To activate protein A-MNase, incubation buffer (3.5mM HEPES, 10mMCaCl_2_, 0.005% digitonin) was added and the DNA was digested for 30min on ice. The incubation buffer was discarded and the reaction was stopped by resuspension in STOP buffer (170 mM NaCl, 20 mM EDTA, 0.005% Digitonin, 50μg/ml RNAseA, 25μg/ml Glycogen). The protein-DNA complex was released by incubation at 37°C for 30min, the supernatant was transferred to a fresh tube and then digested by proteinase K at 50°C for 1 hours. DNA was extracted by ethanol precipitation. Libraries were made with 6 ng of CUT&RUN DNA fragments using the NEBNext Ultra II DNA Library Prep Kit for Illumina (NEB #E7645S). The manufacturer’s protocol was adjusted to account for shorter DNA fragments as described previously (33). Briefly, end prep was performed at 20°C for 30min followed by 50°C for 1h. The adaptor was used at a concentration of 0.5 μM. 15 PCR cycles were performed at the following conditions: Initial denaturation: 98°C for 30sec. Denaturation 98°C for 10sec; Annealing/Extension 65°C for 10sec and final Extension 65°C for 5min.

### Transwell migration assay

MCF10A-YAP5SA and MCF10A-YAP5SA-ΔNP63 cells were starved for 24h in medium supplemented with 0.25% horse serum. Membrane well inlets (OMNILAB) were equilibrated and the top chamber of the transwell was loaded with 500 μl cell suspension (40.000 cells/ml). The lower chamber was filled with 600 μl of MCF10A complete medium. Cells were incubated at 37°C in 5% CO_2_ for 40 h. Transwell inlets were removed and rinsed in PBS and cells on the upper side of the transwell were wiped off with cotton swabs. Migrated cells on the lower side were fixed for 10 minutes in ice-cold methanol and stained with crystal violet 2% in methanol for 20 minutes, followed by three washing steps with 1 x PBS. Migrated cells were photographed by an inverted Leica DMI 6000B microscope. Crystal violet was solubilized by the addition of 33% acetic acid and measured at 595nm in a Multiscan Ascent microtiter plate reader (Labsystems).

### Mammosphere assay

Cells were trypsinized and resuspended in mammosphere medium (DMEM/F12 supplemented with 1% penicillin/streptomycin, 52 μg/ml bovine pituitary extract (Thermo Fisher Scientific), 0.5 μg/ml hydrocortisone (Sigma), 5 μg/ml insulin (Sigma), 100 ng/ml EGF (Sigma) and 1x B27 supplement (Thermo Fisher Scientific). Single-cell suspensions were obtained by resuspending the cells 8 times using a 10 ml syringe (25G needle). Finally, 2000 cells were seeded into the wells of 24-well plates in mammosphere medium. Mammospheres were counted after 7 days.

### siRNA transfection

Double-stranded RNA was purchased from Eurofins or Thermo Fischer Scientific. siRNAs were transfected in a final concentration of 30 nM using RNAiMAX (Thermo Fisher Scientific) according to the manufacturer’s protocol. siRNAs are listed in Supplemental Table 1.

### Lentiviral production and infection

Lentiviral particles were generated in HEK293T co-transfected with psPAX2, pCMV-VSV-G and a lentiviral vector. Filtered viral supernatant was diluted 1:1 with culture medium and supplemented with 4 μg/ml polybrene (Sigma). Infected cells were selected 48 h after infection with 300 μg/ml neomycin for 7 days.

### CRISPRi

MCF10A-YAP5SA cells expressing Cas9-KRAB were generated by infection with lentiviral vector pHAGE TRE dCas9-KRAB (Addgene #50917; (34)) and selection with neomycin. Individual clones were isolated and screened by immunostaining for homogenous nuclear expression of dCas9-KRAB using HA-antibodies. Guide RNAs were designed with the sgRNA designer of the Broad Institute (https://portals.broadinstitute.org) (35). Guide RNAs were first cloned individually into lenti-sgRNA-blast (Addgene #104993) (36) which contains an U6 promoter and a sgRNA scaffold. To express all five guide RNA cassettes from a single lentiviral vector, we created multiplex-lenti-sgRNA-blast by replacing the KpnI-EcoRI fragment of lenti-sgRNA-blast with annealed oligos SG2790 and SG2791. Next, sgRNA-cassettes were amplified by PCR and assembled into multiplex-lenti-sgRNA-blast to generate multiplex-lenti-sgRNA-blast-CDC20. Guide RNA sequences and PCR primer sequences are listed in Supplemental Table 1. MCF10A-YAP5SA-dCas9-KRAB cells were infected with either multiplex-lenti-sgRNA-blast-CDC20or with lenti-sgRNA-blast with a nonspecific control guide RNA (lenti-sgRNA-blast-control) and selected with blasticidine. Guide RNA sequences are listed in Supplemental Table 1.

### RT-qPCR

Total RNA was isolated using peqGOLD TriFast (Peqlab) according to the manufacturer’s instructions. RNA was transcribed using 100 units RevertAid reverse transcriptase (Thermo Fisher Scientific). Quantitative real–time PCR reagents were from Thermo Fisher Scientific and real-time PCR was performed using a Mx3000 (Stratagene) and qTower3G (Analytik Jena). Expression differences were calculated as described before (27). Primer sequences are listed in Supplemental Table 1.

### ChlP-qPCR and ChlP-seq

Cells were cross-linked with 1% formaldehyde (Sigma) for 10 min at room temperature. The reaction was stopped by adding 125 mM glycine (Sigma). After cells were lysed for 10 minutes on ice [5 mM PIPES pH 8.0, 85 mM KCl, 0.5% NP40, 1 mM PMSF, protease inhibitor cocktail (Sigma)], nuclei were resuspended in RIPA buffer [50 mM HEPES pH 7.9, 140 mM NaCl, 1 mM EDTA, 1% Triton X-100, 0.1% sodium deoxycholate, 0.1% SDS, 1 mM PMSF, protease inhibitor cocktail (Sigma)]. Chromatin was fragmented to an approximate length of 150 to 300 bp using a Branson sonifier. Antibodies (3 μg for ChIP-qPCR and 9 μg for ChIP-seq) were coupled to protein G dynabeads (Thermo Fisher Scientific) for 6 hours at 4°C and then incubated with fragmented chromatin over night at 4°C. Beads were washed in total twelve times with wash buffer I (50mM Tris-HCl pH8, 0.15M NaCl, 1mM EDTA, 0.1% SDS, 1% Triton X-100, 0.1% sodium deoxycholate), wash buffer II (50mM Tris-HCl pH8, 0.5M NaCl, 1mM EDTA, 0.1% SDS, 1% Triton X-100, 0.1% sodium deoxycholate), wash buffer III (50mM Tris-HCl pH8, 0.5M LiCl_2_, 1mM EDTA, 1% Nonidet P-40, 0.7% sodium deoxycholate) and wash buffer IV (10mM Tris-HCl pH8, 1mM EDTA). 1mM PMSF and protease inhibitor cocktail (Sigma) was added freshly to all buffers. Chromatin was eluted in (10mM Tris-HCl pH8, 0.3M NaCl, 5mM EDTA, 0.5% SDS, 10μg/ml RNAseA) and crosslink was reversed at 65°C over night. Proteins were digested by adding 200 μg/ml proteinase K at 55°C for 2 hours. DNA was purified using the QIAquick PCR Purification Kit (QIAGEN) and eluted in 50 μl EB buffer. ChIP samples were analyzed by qPCR or subjected to library preparation according to the manufacturer’s protocol (NEBNext Ultra II DNA Library Prep Kit for Illumina, NEB) using Dual Index Primers (NEBNext Multiplex Oligos for Illumina, NEB). Libraries were sequenced on the NextSeq 500 platform (Illumina). Antibodies used for ChIP are listed in Supplemental Table 2.

### Immunoblotting and immunoprecipitation

Cells were lysed in TNN [50 mM Tris (pH 7.5), 120 mM NaCl, 5 mM EDTA, 0.5% NP40, 10 mM Na_4_P_2_O_7_, 2 mM Na_3_VO_4_, 100 mM NaF, 1 mM PMSF, 1 mM DTT, 10 mM ß-glycerophosphate, protease inhibitor cocktail (Sigma)]. Proteins were separated by SDS-PAGE, transferred to PVDF membrane and detected by immunoblotting.

For co-immunoprecipitation of endogenous proteins from nuclear extracts, cells were first lysed in [10 mM HEPES pH 7.4, 10 mM NaCl, 3 mM MgCl_2_, protease inhibitor cocktail (Sigma)] for 20 min on ice. Pelleted nuclei were resuspended in nuclei lysis buffer [20 mM HEPES pH 7.4, 400 mM NaCl, 1.5 mM MgCl_2_, 0.1 mM EDTA, 15% glycerol, 0.5 mM DTT, protease inhibitor cocktail (Sigma)] mixed 1:1 with TNN buffer for another 20 min on ice. After spinning at full speed for 10 min, the supernatant was mixed with 20 mM HEPES pH 7.4 and used for immunoprecipitation overnight at 4°C in constant rotation. Complexes were collected with protein G-dynabeads (Thermo Fisher Scientific) for 2h at 4°C. Beads were washed three times with nuclei lysis buffer mixed 1:1 with 20 mM HEPES pH 7.4. Proteins were eluted from beads and bound proteins were detected by immunoblotting. Antibodies are listed in Supplemental Table 2.

### Immunostaining

Cells were fixed with 3% paraformaldehyde and 2% sucrose in PBS for 10 minutes at room temperature. Cells were permeabilized using 0.2% Triton X-100 (Sigma) in PBS for 5 minutes and blocked with 3% BSA in PBS-T (0.1% Triton X-100 in PBS) for 30 minutes. Primary antibodies were diluted in 3% BSA in PBS-T and incubated with the cells for 1 hour at room temperature or overnight for cell cycle profile. After three washing steps with PBS-T, secondary antibodies conjugated to Alexa 488 and 594 (Thermo Fisher Scientific) and Hoechst 33258 (Sigma) were diluted 1:500 in 3% BSA in PBS-T and incubated with the coverslips for 30-60 minutes at room temperature. Finally, slides were washed three times with PBS-T and mounted with Immu-Mount™ (Thermo Fisher Scientific). Pictures were taken with an inverted Leica DMI 6000B microscope equipped with a Prior Lumen 200 fluorescence light source and a Leica DFC350 FX digital camera.

### PLA

PLA was performed using the Duolink In Situ Kit (Sigma) according to the manufacturer’s instructions. The following antibodies were used: LIN9 (Bethyl, A300-BL2981) and CDK7 (Cell Signaling 2912). Pictures were taken with an inverted Leica DMI 6000B microscope equipped with a Prior Lumen 200 fluorescence light source and a Leica DFC350 FX digital camera.

### Bioinformatics

Base calling was performed with Illumina’s CASAVA software or FASTQ Generation software v1.0.0 and overall sequencing quality was tested using the FastQC script. Read files were imported to the Galaxy Web-based analysis portal (37). Within Galaxy, ATC-seq and ChIP-seq reads were mapped to the human genome (hg19 assembly) using Bowtie2 with default parameters (38). For ATAC-seq properly paired reads were filtered using a Phred score cutoff of 30 and mitochondrial reads were removed. ATACseq peaks were called using Genrich (https://github.com/jsh58/Genrich). Differential peak analysis was performed within Galaxy with Limma-voom (39). Nucleosome calling was carried out with NucleoATAC (40). To create heat maps and density profiles of ATACseq, ChIPseq and CUT&RUN data, DeepTools 2 was used (41). First, normalized bigWig files were created using bamcoverage with a bin size of 10 and normalizing to counts per million (CPM). BigWig files were used to compute reads centered on ATAC-seq peak summits (called with Genrich), YAP peak summits or on the TSS of MMB-target genes (42) using computeMatrix. Heatmaps and profiles were created with the plotHeatmap and plotProfile tools. Metagene plots of PolII enrichment across MMB target genes were created with plotProfiles. ChIPseq and ATACseq data were visualized with the Integrated Genome Viewer (43). For ChIP-seq of H3K4me1, H3K4me3, H3K27ac, B-MYB and LIN9 in control MCF10A cells and YAP5SA cells, we reanalyzed or previously published datasets which are available under GEO accession number GSE115787 (21)

## QUANTIFICATION AND STATISTICAL ANALYSIS

Statistical analyses were performed using Prism 9 (GraphPad). Tests used to determine statistical significance are indicated in the figure legends. Comparison of two groups was done by a two-sided Student’s t test. *P*-values <0.05 were considered statistically significant. * *P* <0.05;** *P* <0.01; *** *P* <0.001; **** *P* <0.0001.

## RESULTS

### YAP5SA results in widespread changes in chromatin accessibility

To recapitulate events in YAP induced tumorigenesis we used untransformed MCF10A cells stably expressing doxycycline-inducible YAP5SA, a constitutive active allele of YAP that cannot be inhibited by the Hippo kinases. In this system, YAP5SA is robustly induced by doxycycline treatment for 48 hours (Figure 1A). To determine whether YAP leads to changes in chromatin accessibility, we performed *assay for transposase-accessible chromatin with sequencing* (ATAC-seq). We observed that the expression of YAP5SA resulted in widespread changes in chromatin accessibility in MCF10A cells (Figures 1A,B). Overall, we identified 23,890 newly accessible ATAC-seq peaks “opened” and 13,612 less accessible peaks “closed” in YAP5SA expressing cells compared to control cells. In contrast, 110,403 “flat” peaks were accessible in both conditions and did not change upon expression of YAP5SA. The vast majority of opened and closed peaks mapped to intergenic and intronic regions and only very few peaks were found in promoter regions of annotated genes (Figure 1C).

**Figure 1:**
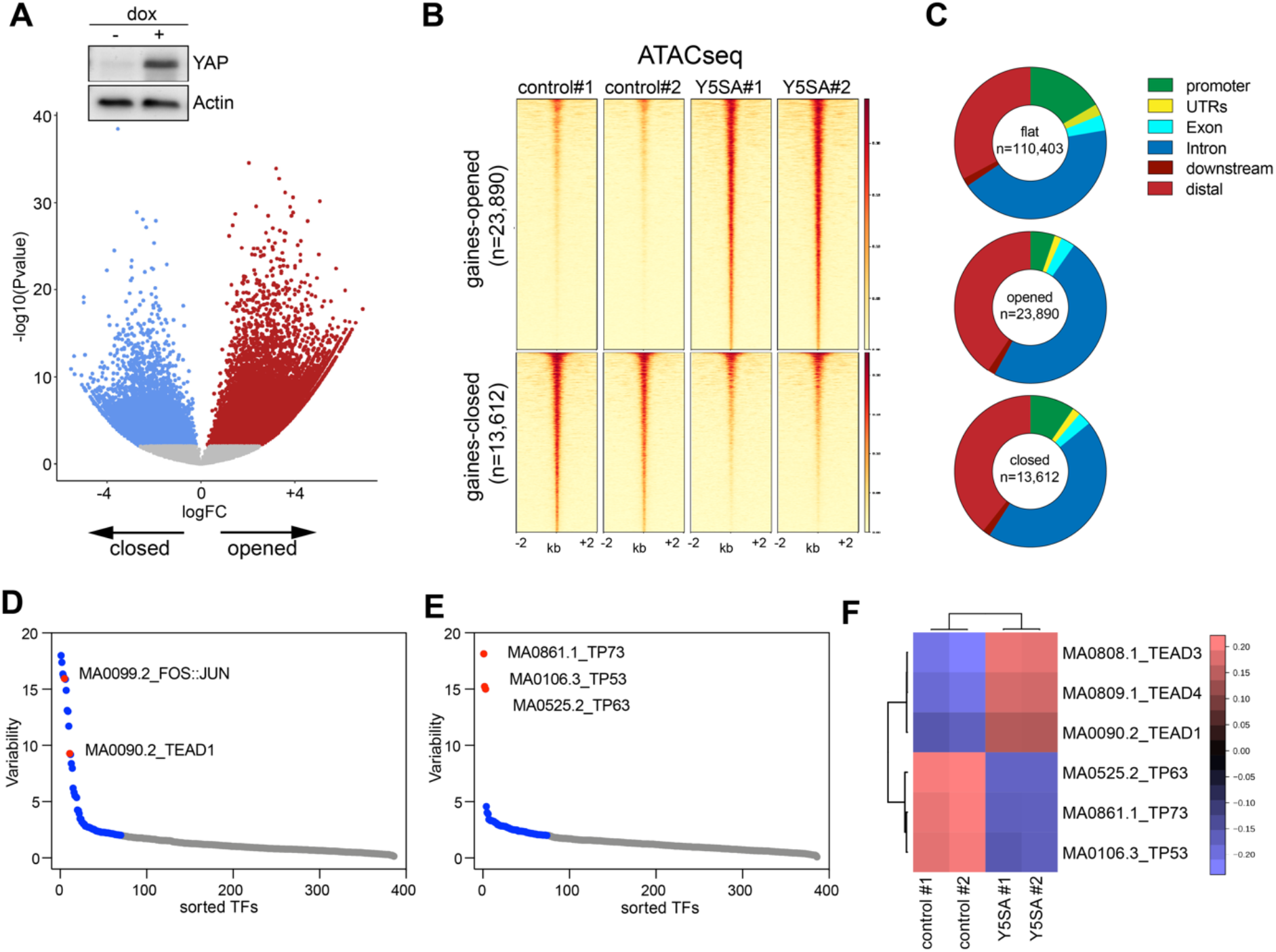
Expression of YAP5SA results in genome-wide changes in chromatin accessibility. A) Volcano plot of ATAC-seq data after induction of YAP5SA in MCF10A cells expressing doxycycline-inducible YAP5SA. 23,890 regions were newly opened, 13,612 newly closed, 110,403 were unchanged (flat) at q<0.035. 2 biological replicates per condition. The insert shows YAP expression as analyzed by immunoblotting. ß-Actin served as a control. dox= doxycycline B) Heatmap of upregulated and downregulated ATAC-seq peaks in a window of −2 kb to + 2 kb centered on the middle of the peak. C) Distribution of ATAC-seq peaks relative to known genes in the genome. D) and E) ChromVAR chromatin variability scores for ATAC-seq data of YAP5SA expressing MCF10A cells, indicating TEAD and p53-family binding sites as the most variable motifs in gained open (D) and gained closed regions (E), respectively. F) Heatmap showing motif enrichment for open and closed regions.

For an unbiased identification of transcription factor motifs associated with regions with differential chromatin accessibility following expression of YAP5SA we used chromVAR (44). The top motifs enriched in chromatin regions that are gained accessible after expression of YAP5SA correspond to the binding sites for TEAD proteins and for the Activator Protein-1 (AP-1) family of transcription factors (Figure 1D, F). This is consistent with the previous finding that YAP is recruited to the chromatin by TEAD proteins and that TEAD and AP-1 interact at enhancers to drive the expression of YAP-dependent genes (17). Interestingly, binding motifs for the p53 family of transcription factors were highly enriched in regions that became less accessible in YAP5SA expressing cells, suggesting that the p53 family, comprised of the three members p53, p63, and p73, could play a role in shaping the YAP-mediated enhancer landscape (Figure 1E, F) (see below).

### YAP5SA invades a subset of enhancers leading to their opening and hyperactivation

We first focused on intergenic regions that become more accessible in cells expressing YAP5SA. We used our previous ChIP-seq data sets to identify primed and active enhancers in MCF10A cells. Enhancers were defined as H3K4me1-positve/ H3K4me3-negative regions that are not within 1 kb of a transcription start site (Figure 2A). By this approach we identified a total of 34,469 putative enhancers in control cells and YAP5SA expressing cells. We clustered these enhancers in two categories based on accessibility following YAP5SA expression. We found that previously accessible enhancers are further opened upon expression of YAP5SA (Figure 2B). Opened enhancer regions also become activated as indicated by increased levels of H3K27ac, which was used as an established indicator of enhancer activity.

**Figure 2:**
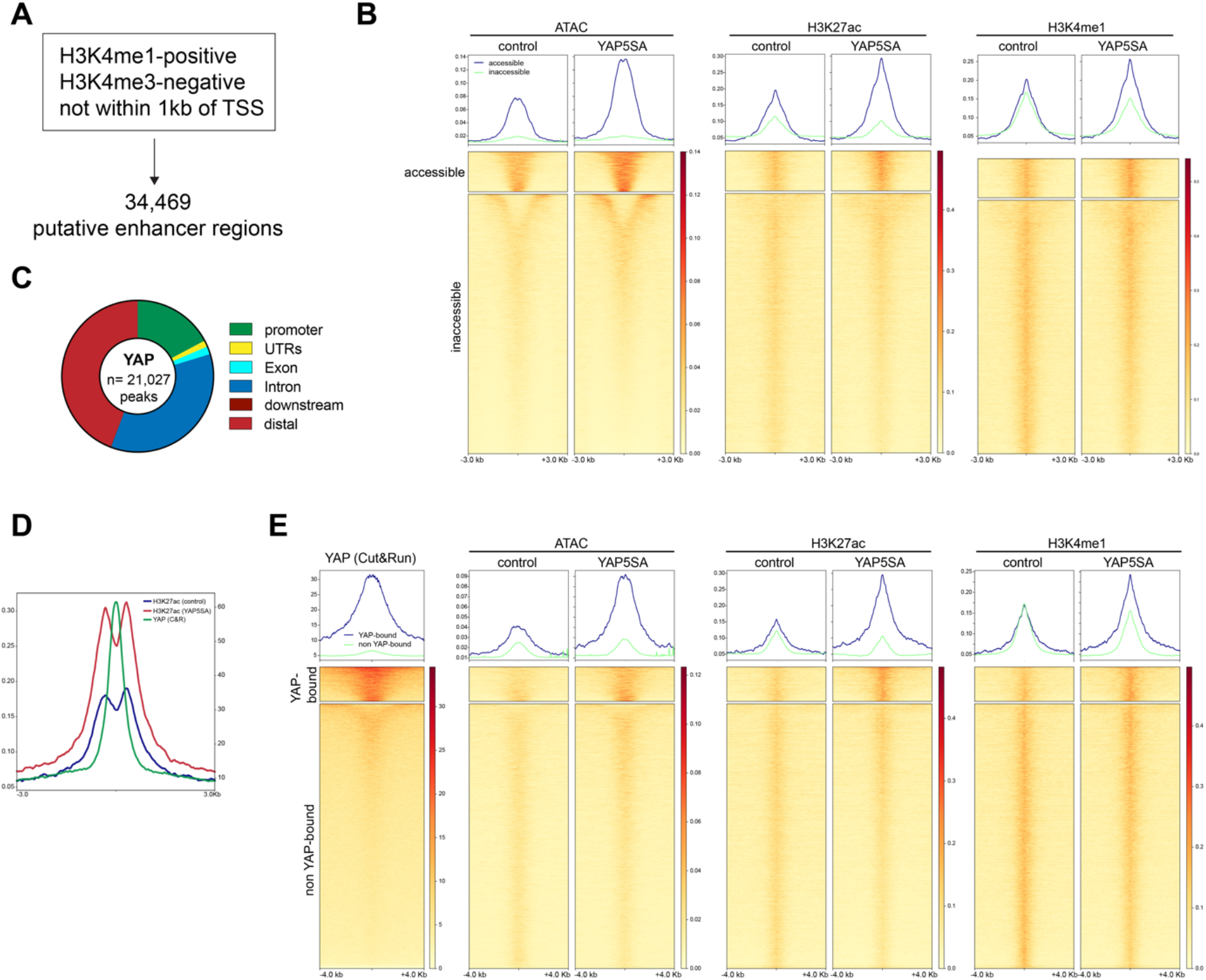
YAP5SA invades a subset of enhancers leading to their opening and hyperactivation. A) Identification of 34,469 putative enhancer regions by analysis of H3K4me1 and H3K4me3 ChIPseq data. B) Line profiles and heatmap displaying ATACseq and ChIPseq. k-means clustering was performed according to ATAC-seq data. All data sets are arranged to match the order of enhancers found by clustering according to accessibility. C) Binding sites for YAP5SA in MCF10A cells as determined by Cut&Run. The location of YAP peaks relative in relation to genomic features is shown. D) Line plot of enrichment of YAP and H3K27Ac at YAP enhancer binding sites showing that the YAP-peaks are not randomly distributed in relation to H3K27ac but that the center of the YAP peak is flanked by two H3K27ac peaks that gain acetylation when YAP5SA is expressed. E) Line profiles and heatmap displaying YAP Cut&Run, ATACseq and ChIPseq data. k-means clustering according to YAP enrichment at the enhancer regions identified YAP-bound and non-YAP bound enhancers. All data sets are arranged to match the order of enhancers found by clustering according to YAP enrichment.

We next profiled YAP by *Cleavage Under Targets and Release Using Nuclease* (CUT&RUN), which has a better signal to noise ratio compared to ChIP-seq and is particularly suited for factors that do not directly bind to DNA (31). We identified 21,027 high confidence YAP peaks compared to the 5,630 peaks previously identified by ChIP-seq in MCF10A cells, confirming the higher sensitivity of the Cut&Run method. YAP5SA binds mostly to distal and intergenic regions, consistent with previous data (Figure 2C). Plotting the enrichment of H3K27Ac at YAP enhancer binding sites shows that the YAP peaks are not randomly distributed in relation to H3K27ac but that the center of the YAP peak is flanked by two H3K27ac peaks that gain acetylation when YAP5SA is expressed (Figure 2D).

Clustering of primed and active enhancers based on YAP enrichment by CUT&RUN revealed that accessibility at YAP-bound regions increased upon expression of YAP5SA, while it did not change at the non-YAP-bound regions. Notably, baseline accessibility at YAP-bound enhancers in control cells was higher compared to non-YAP-bound regions, suggesting that YAP preferentially binds to regions that are partially accessible but no to completely closed chromatin. YAP-binding not only increased enhancer accessibility but also resulted in enhancer hyper-activation based on the H3K27ac ChIP-seq signal. In contrast, non-YAP-bound enhancers were not further activated by YAP5SA. Collectively, these data suggest that YAP5SA invades a significant subset of all enhancers in MCF10A cells leading to their opening and hyperactivation.

### YAP5S does not increase chromatin accessibility at MMB-regulated promoters

To explore how YAP5SA-activated enhancers control gene expression, we focused on cell cycle genes co-regulated by YAP and by the Myb-MuvB (MMB) complex. We have previously shown that pro-tumorigenic functions of YAP, such as cell cycle entry and mammosphere formation depend on activation of MMB by YAP (21). Furthermore, the expression of genes coactivated by YAP and B-MYB is associated with poor survival of cancer patients. By binding to distant enhancers YAP stimulates the association of the B-MYB subunit of MMB to promoters of MMB-target genes, providing an explanation for their increased expression (21). Plotting the ATAC-seq signal at MMB target genes showed that the TSS is accessible in control cells, although nucleosome occupancy at the −1 nucleosome was slightly reduced when YAP5SA was expressed (Supplemental Figure S1A). In contrast, at the TSS of 1233 genes with gained ATAC-seq peaks in the promoter (see Figure 1C), accessibility increased and nucleosome occupancy decreased (Supplemental Figure S1B). Nucleosome positioning and nucleosome occupancy at the TSS of MMB-target genes and at genes with gained ATAC-seq peaks did not change when YAP5SA was expressed (Supplemental Figure S1A,B). Plotting the ChIP-seq signal for LIN9, a subunit of the MuvB core, revealed that LIN9 is present at the TSS of MMB target genes, but not at genes which opened ATAC-seq regions. Notably. LIN9 was present at the TSS of MMB-target genes before they were activated by YAP5SA, providing a possible explanation for the constitutive accessibility of the TSS. The accessible region at the TSS of MMB target genes also overlaps with the binding sites for B-MYB and FOXM1, which are recruited to the TSS following expression of YAP5SA.

Overall, these data suggest that activation of MMB-target genes is regulated through a different mechanism rather than opening and remodeling of the chromatin at the TSS of these genes.

### YAP regulated enhancers facilitate RNA pol II Ser5 phosphorylation at the CDC20 locus

We next investigated regulation of CDC20 as an example of a YAP/MMB co-regulated gene. YAP activates CDC20 expression by binding to two distal enhancers that interact with the CDC20 promoter by chromatin looping (Pattschull et al., 2019 and Supplemental Figure S1C). Accessibility and acetylation of H3K27ac at these two enhancers increased in cells expressing YAP5SA (Supplemental Figure S1C,D). To better understand how the YAP-bound enhancers control the expression of CDC20, we inactivated the enhancers by CRISPR interference (CRISPRi), a CRISPR/Cas9 epigenetic tool based on catalytically-inactive dCas9 fused to the KRAB transcriptional repressor domain (dCas9-KRAB) (Figure 3A). We created MCF10A-YAP5SA cells stably expressing doxycycline-inducible dCas9-KRAB together with either a nonspecific control guide RNA or a set of five guide RNAs that target dCas9-KRAB to the two CDC20 enhancers (Figure 3B). Western blotting and immunostaining confirmed doxycycline-dependent expression of dCas9-KRAB and YAP5SA (Supplemental Figure S2). To investigate whether dCas-Cas9 prevents enhancer activation, we performed ChIP assays with antibodies specific for acetylated H3K9, H3K27, H4 and H2A.Z, chromatin marks that are associated with transcriptional activation. YAP5SA resulted in increased histone acetylation at the two YAP-bound enhancers, which was prevented when Cas9-KRAB was co-expressed with enhancer-specific guide RNAs (Figure 3C). This not only confirms that the CDC20 enhancers are activated when YAP5SA is expressed but also shows that Cas9-KRAB targeted to the enhancers interferes with YAP-induced enhancer activation by preventing histone acetylation. Importantly, YAP-mediated induction of *CDC20* mRNA expression was abolished when the enhancers were epigenetically silenced, indicating that the identified enhancers are required for YAP5SA-mediated expression of CDC20 (Figure 3D). As a control, silencing of the CDC20 enhancers did not affect induction of *AMOTL2* by YAP5SA, a gene regulated by binding of YAP to the promoter.

**Figure 3:**
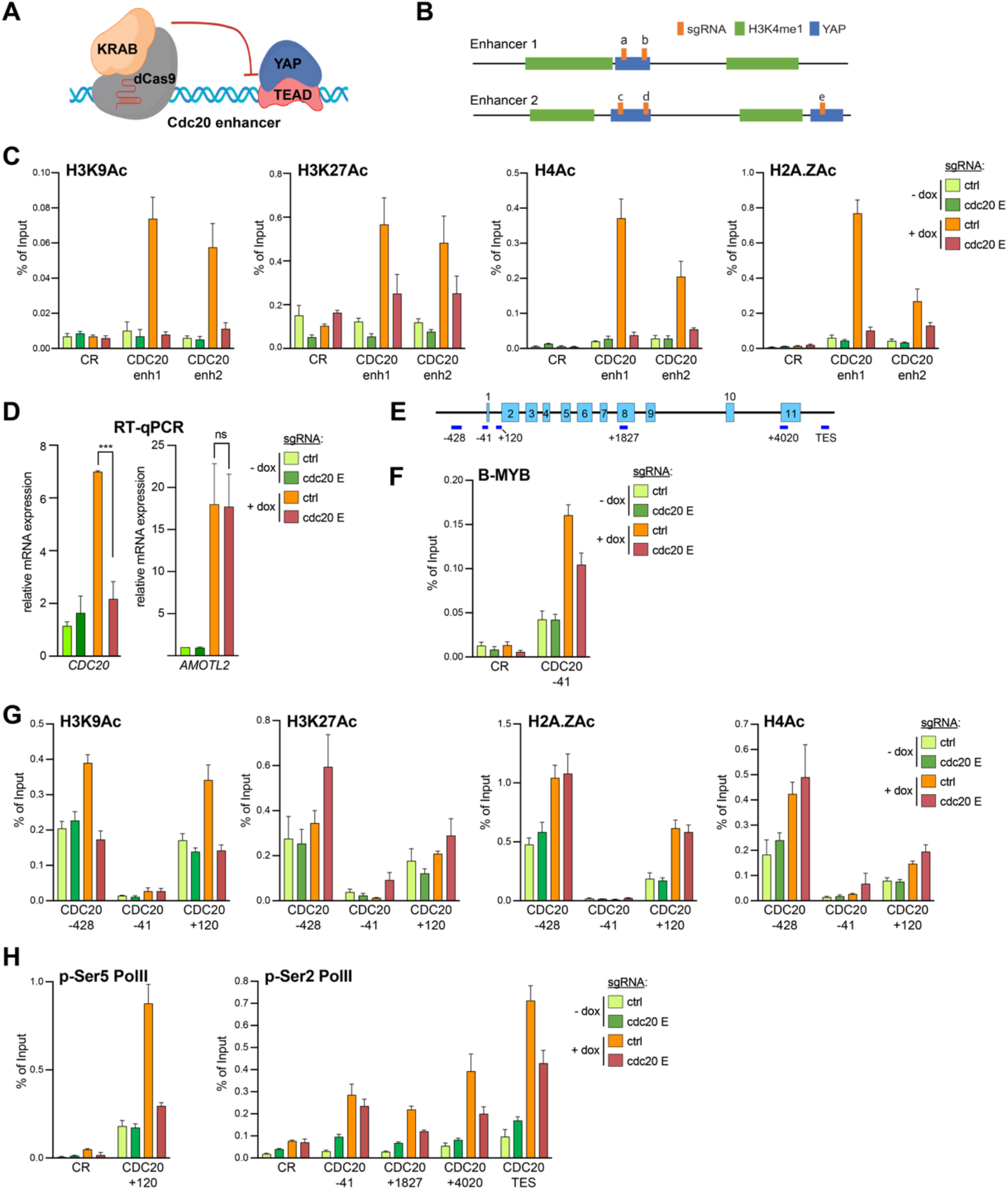
A role for YAP-bound enhancers in histone acetylation and RNA Pol II Ser5 phosphorylation at the CDC20 locus. A) Illustration of the CRISPR-interference (CRSPRi) system to inhibit YAP-bound enhancers. B) Scheme depicting the tow CDC20 enhancers (E1 and E2) and position of sgRNAs (a-e) in relation to the YAP (blue) and H3K4me1 (green) peaks as determined by ChIPseq. C) ChIP-qPCR for H3K9Ac, H3K27Ac, H4Ac and H2A.Zac at the two CDC20 enhancers before and after YAP5SA induction in cells expressing either a control guide RNA or enhancer-specific guide RNAs demonstrating that targeted CRISPRi interferes with enhancer activation by YAP5SA. CR: negative control region. D) MCF10A-YAP5SA-Cas9-KRAB cells expressing either control or enhancer-specific guide RNAs were treated with doxycycline (+dox) to induce the expression of YAP5SA and Cas9-KRAB or were left untreated (-dox). The expression of *CDC20* and *AMOTL2* relative to *GAPDH* was analyzed by RT-qPCR. Error bars: represent SD. N=3 independent replicates. Student’s t-test. **** = p<0.0001, ns = not significant. E) Scheme of the CDC20 locus and the position of amplicons used for ChIP-qPCR. F, F-H) ChIP-qPCRs at CDC20 indicated locus for F) B-MYB, for G) H3K9Ac, H3K27Ac, H4Ac, and H2A.Zac for H) p-Ser5 Pol ll, p-Ser2 Pol ll and before and after YAP5SA induction in MCF10A-Cas9-KRAB cells expressing either a control guide RNA or enhancer-specific guide RNAs. CR: negative control region. For all ChIP-qPCR assays the mean and SDs of technical triplicates of a representative experiment (n=2 to 3 biological replicates) are shown.

We next explored how silencing of the enhancers affects the CDC20 locus (Figure 3E). By ChIP-qPCR, YAP5SA increased the binding of B-MYB to the CDC20 TSS, as previously shown (Figure 3F). Silencing the enhancers reduced binding of B-MYB, but did not completely prevent it. This suggests that enhancer-activation only partially controls the recruitment of B-MYB, consistent with the dual role of YAP in promoting the chromatin binding of B-MYB as well as increasing the mRNA expression of *MYBL2* (21).

To further investigate the mechanism by which enhancer activation results in CDC20 promoter activation, we investigated acetylation of H2A.Z, H3K9, H3K27 and H4 by ChIP (Figure 3G). Acetylated histones showed the expected bimodal distribution around the transcriptional start site of CDC20. The signal for H2A.Zac, H4ac and H3K9ac was increased in cells expressing YAP5SA. Notably, only the increase in acetylation of H3K9 was dependent on enhancer activity while acetylation of H2A.Z and H4 was not prevented and H3K27 acetylation was increased by disruption of enhancer activation. Enhanced acetylation of H4 has been reported before after inhibition of CDK7, the kinase that is responsible for phosphorylating RNA Pol II at Ser5 (45). Similarly, depletion of ARID1A, which regulates promoter proximal pausing leads to increased H3K27 acetylation at the +1 nucleosome (46). Taken together these data suggest that YAP5SA-mediated CDC20 enhancer activation has a more direct impact on H3K9-acetylation at the CDC20 promoter than on acetylation of H4, H2A.Z and H3K27.

To further investigate the functional role of the YAP-bound enhancers on RNA Pol II dynamics, we next focused on the recruitment of RNA Pol II phosphorylated at Ser5 (p-Ser5), which is associated with the transition from initiation to elongation and phospho-Ser2 (p-Ser2), which occurs during pause release and is associated with the elongating RNA Pol II throughout gene bodies. As YAP has primarily been implicated in stimulating transcriptional elongation through controlling the pause-release step (19), we expected an increase in the levels of p-Ser2 and reduced Ser5-phosphorylation of Pol II in cells expressing YAP5SA and possibly accumulation of p-Ser5 Pol II upon enhancer-inhibition. Instead, the expression of YAP5SA caused a robust increase in p-Ser5 Pol II at the TSS of CDC20 and, importantly, this increase was prevented by CRISPRi-mediated enhancer inhibition (Figure 3H). YAP5SA also caused an enrichment of RNA Pol II p-Ser2 in the CDC20 gene body and at the TES, which was partially rescued by CRISPRi (Figure 3H). Taken together these observations suggest that enhancer inhibition prevents accumulation of paused Pol II at the CDC20 promoter by YAP5SA. Inhibition of the enhancer also reduced Pol II Ser2 phosphorylation and transcriptional elongation, but this might be indirect to the effect on initiation and promoter escape.

To assess changes in Pol II occupancy not only at the CDC20 gene but at a larger panel of MMB target genes, we measured genome-wide occupancy of Pol II phosphorylated at Ser5 by ChIP-seq. RNA Pol II p-Ser5 was increased at the TSS-proximal regions of MMB-targets after YAP5SA expression. Using ChIP-seq and an antibody that primarily recognizes unphosphorylated Pol II, we found that levels of unphosphorylated Pol II were also increased at MMB targets by YAP5SA albeit to a lesser extent compared to Ser5 phosphorylated Pol II (Figure 4A). ChIP-seq for RNA Pol II phosphorylated at Ser2 revealed higher levels of p-Ser2 Pol II in gene bodies and at the TES of MMB targets after expression of YAP5SA, consistent with previous studies that linked YAP to transcriptional elongation. Plotting the fold enrichment of unphosphorylated Pol II, p-Ser5 Pol II and p-Ser2 Pol II showed that p-Ser5 boasted the biggest increase in cells expressing YAP5SA (Figure 4B). Genome browser tracks of the MMB target genes CCNF, TOP2A, NEK2 and CDC20 illustrating these findings are shown in Figure 4C. Taken together these observations suggest that YAP stabilizes the initiating or paused RNA Pol II at MMB-target genes, extending previously published studies that suggested YAP primarily regulates the pause-release step and transcriptional elongation.

**Figure 4:**
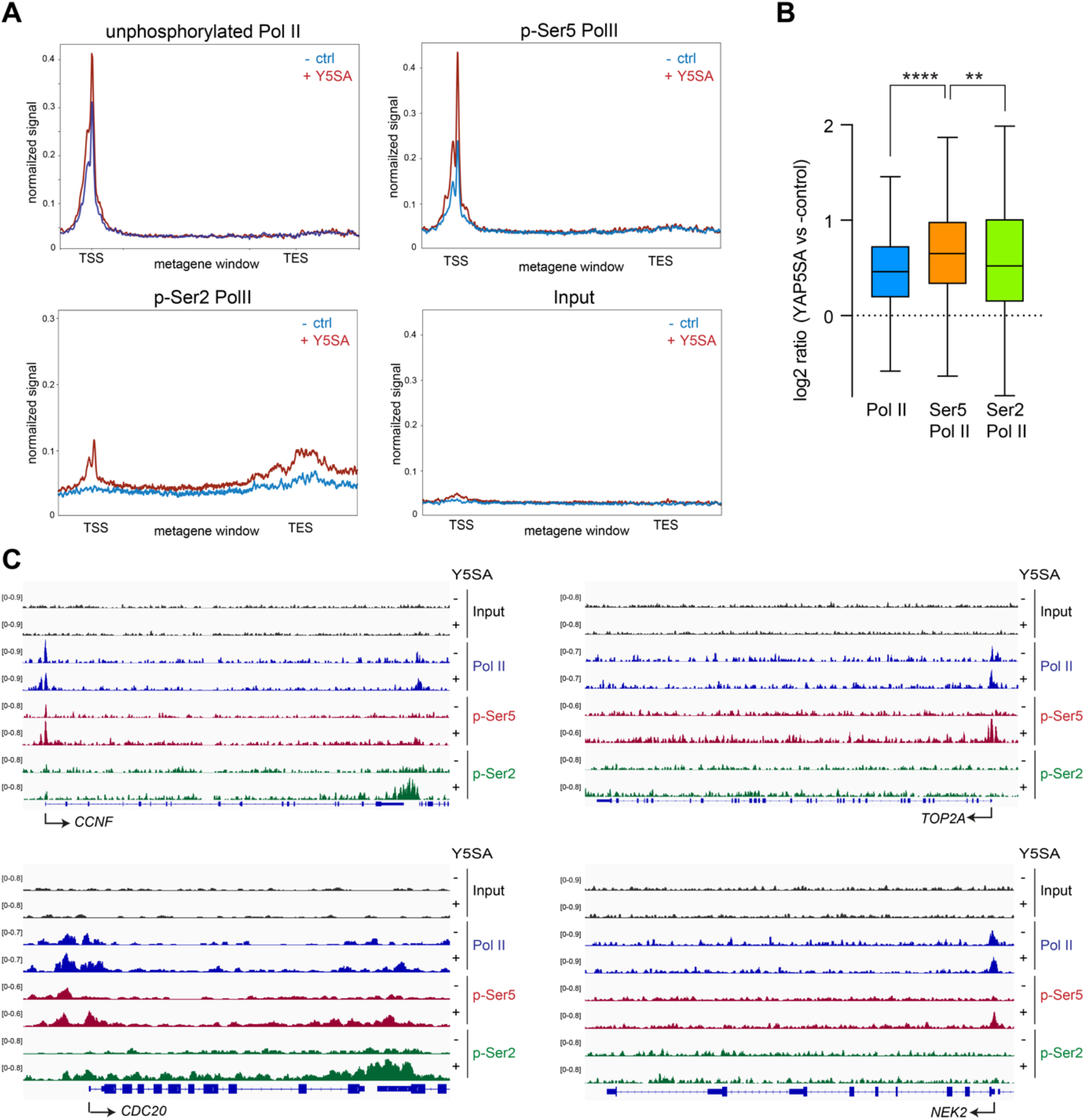
Enrichment of Pol II phosphorylated at Ser5 at MMB-target genes in cells expressing YAP5SA. A) Metagene plots of unphosphorylated RNA Pol II, RNA Pol II phosphorylated at serin 5 (p-Ser5 Polll) or phosphorylated at serine 2 (p-Ser2 Polll) at YAP-MMB-regulated genes. B) Boxplot representing the fold enrichment of unphosphorylated Pol ll and p-Ser5 Pol ll in a window of −500bp to +500bp at the TSS or of p-Ser2 Pol lI in a window of TSS +500 to TES +3000 bp at MMB-regulated genes in cells expressing YAP5SA vs control cells. Student’s t-test. ** = p<0.01, **** = p<0.0001. C) Genome browser ChIP-seq tracks of CCNF, TOP2A, NEK2 and CDC20 demonstrating enhanced phosphorylation of Ser5 Pol II at the TSS by YAP5SA.

### A role for CDK7 in YAP-mediated activation of MMB-target genes

Given that YAP5SA induces Ser5 Pol II phosphorylation at MMB-target genes, we next asked whether there is a functional link between CDK7, the main kinase that phosphorylate RNA Pol II and the expression of these genes by using TZH1 a small molecule CDK7 kinase inhibitor. To exclude indirect effects due to long CDK inhibition, we used ATR-CHK1 pathway inhibition as a tool to achieve rapid activation of MMB-target genes. It has previously been shown that the ATR-CHK1 pathway limits the activity of MMB and FOXM1 during S-phase to prevent expression of mitotic genes in cells that are still replicating their DNA (47, 48). We confirmed that inhibition of CHK1 by prexasertib treatment of MDA-MB-231 cells released from a G1/S block leads to hyperactivation of MMB-target genes CDC20 and AURKA between 2 and 6 hours after the release (Figure 5A). Importantly, co-treatment of cells released from a G1/S block with verteporfin, a drug that disrupts the YAP-TEAD interaction (49), prevented the prexasertib-mediated induction of MMB-targets in S-phase, indicating that this process is YAP-dependent (Figure 5A). Inhibition of CDK7 also abolished prexasertib-mediated hyperactivation of MMB targets, revealing a role for CDK7 in this process (Figure 5B). Although CDK7 has recently been reported to stabilize YAP (50), YAP levels were not affected in our experimental system by short-term inhibition of CDK7 (Supplemental Figure S3A). Thus, reduced levels of YAP do not account for the lower induction of MMB target genes when CDK7 is inhibited. Although co-immunoprecipitation experiments showed no detectable biochemical interaction between CDK7 and B-MYB (Supplemental Figure S3B), we reasoned that YAP could enhance the proximity of CDK7 and MMB. To address this possibility, we performed proximity ligation assays (PLA) using antibodies directed at CDK7 and the LIN9 subunit of MMB. While the single antibodies alone yielded only background levels of fluorescence, specific nuclear interactions were detected when antibodies against LIN9 and CDK7 were used (Figure 5C,D). Importantly, the proximity between CDK7 and LIN9 was strongly enhanced when YAP5SA was induced. Consistent with these data binding of CDK7 to the CDC20 promoter was increased in YAP5SA expressing cells and this was abolished by CRISPRi mediated silencing of the CDC20 enhancers (Figure 5E). Taken together these data suggest a role for the YAP-bound enhancers in the recruitment of CDK7 to MMB-regulated promoters and in the subsequent phosphorylation of Pol II at Ser5.

**Figure 5:**
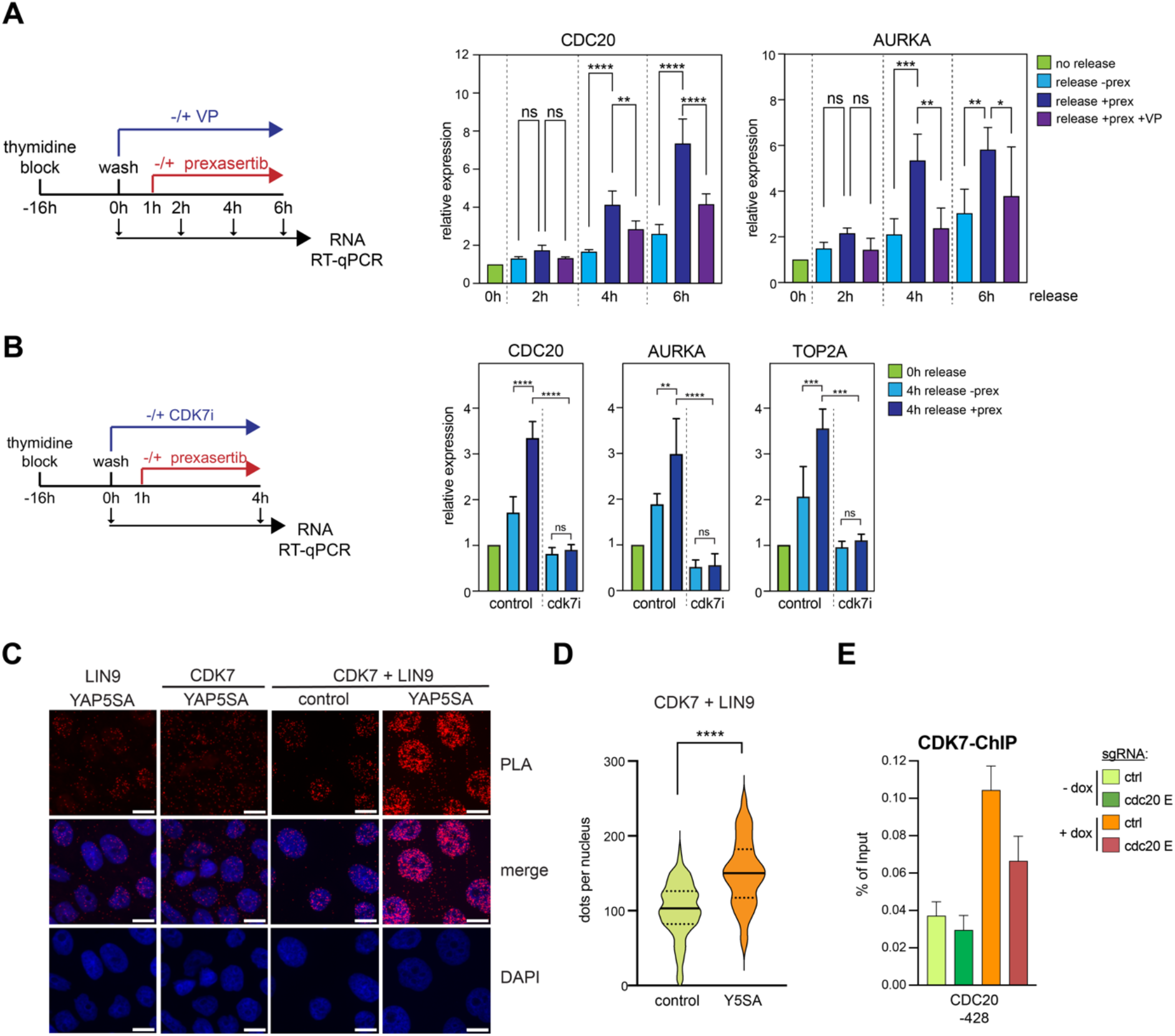
A role for CDK7 in YAP-mediated activation of MMB-target genes. A) Scheme and results of the synchronization experiment with MDA-MB-231 released from a single thymidine block in the presence or absence of verteporfin. One hour after the release, cells were treated with prexasertib or left untreated. 2, 4 and 6 hours after the release, RNA was isolated and subjected to RT-qPCR. B) MDA-MB-231 were released from a single thymidine block in the presence or absence of CDK7i. One hour after the release, cells were treated with prexasertib or left untreated. 4 hours after the release, RNA was isolated and subjected to RT-qPCR. C) Proximity ligation assays (PLA) of YAP and CDK7 and of LIN9 and CDK7 in MCF10A-YAP5SA cells treated with and without doxycycline. Scale bar: 150 μm. D) Quantification of PLA shown in C (n=2 independent experiments). E) ChIP-qPCRs CDK7 binding to the CDC20 locus before and after YAP5SA induction in MCF10A-Cas9-KRAB cells expressing either a control guide RNA or CDC20-enhancer specific guide RNAs. Mean and SDs of technical replicates of a representative experiment (n=3).

### YAP5SA triggers the loss of ΔNp63 from enhancers resulting in reduced chromatin accessibility

As described above, binding motifs for the p53 family of transcription factors were highly enriched in regions that became less accessible in YAP5SA expressing cells (see Figure 1E, F). p53 is expressed at low levels in unstressed cells and because MCF10A cells are known to express p63 but not p73 (51,52), we next focused on p63 as a possible mediator of the reduced chromatin-accessibility following expression of YAP5SA. By ChIP-seq we observed a strong overall reduction in chromatin-binding of p63 after expression of YAP5SA compared to control cells (Figure 6A). Comparison with ATAC-seq revealed that 916 of the 1213 identified high confidence p63 binding sites (q-value < 0.01) were in open chromatin regions in control cells and about 50% of those became inaccessible after YAP5SA expression (Figure 6B). Loss of p63 binding correlated with reduced accessibility, suggesting that p63-binding is required to keep the chromatin accessible (Figure 6C). We next used our previous ChIP-seq data of histone modifications of control and YAP5SA expressing MCF10A cells to determine whether YAP5SA changes the chromatin status at p63 sites. We observed a decrease in H3K27 acetylation at p63 sites, suggesting reduced p63-dependent enhancer activity upon YAP5SA expression (Figure 6D).

**Figure 6:**
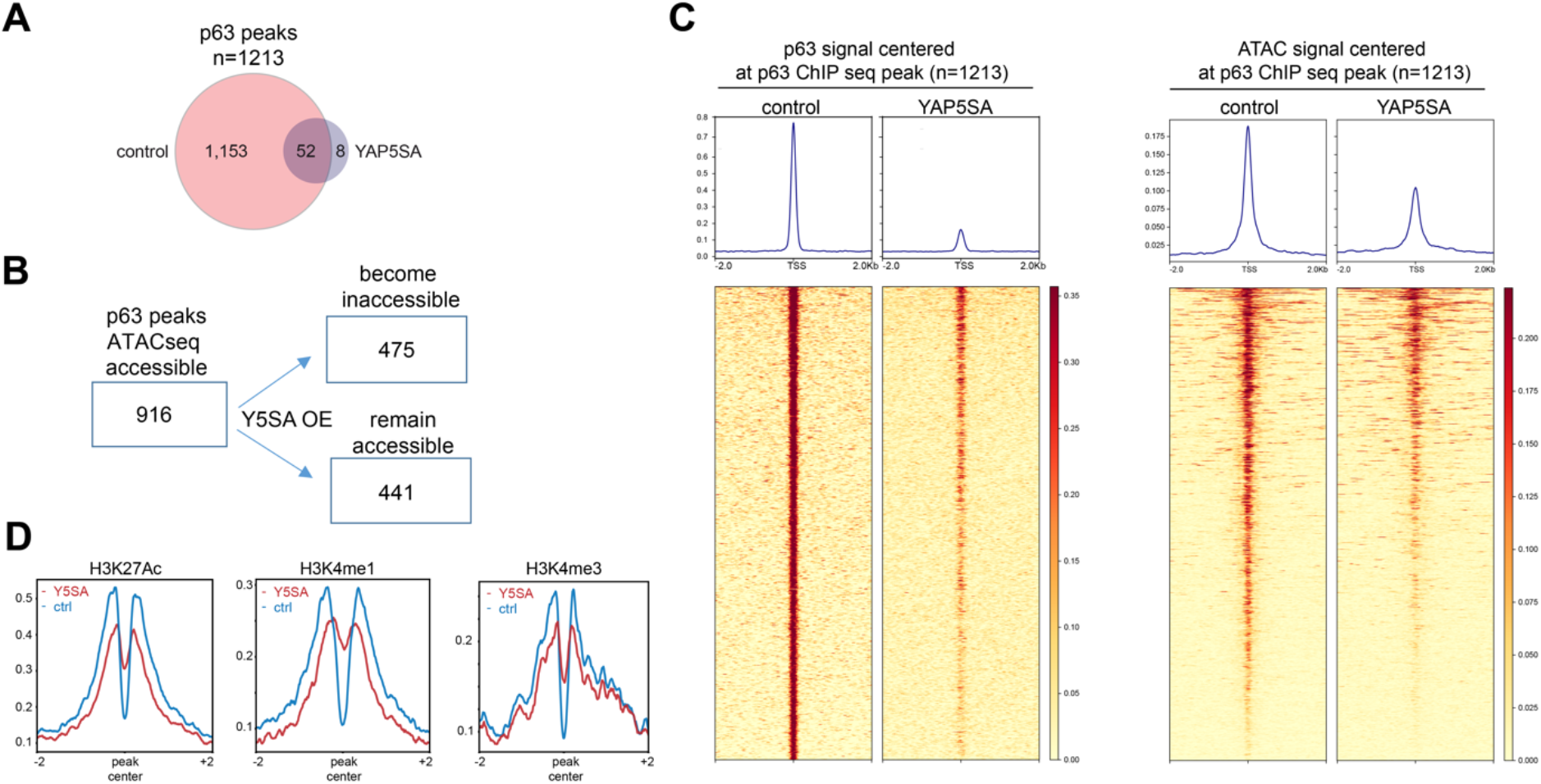
YAP5SA overexpression reduces the chromatin accessibility at ΔNp63 binding sites. A) The genome wide localization of p63 was determined by ChIP-seq with p63 antibodies in MCF10A cells before and after expression of YAP5SA. The number of peaks identified in the two conditions is shown B) Comparison of p63 binding sites with chromatin accessibility obtained by ATAC-seq after the expression of YAP5SA. C) Heatmaps showing p63 enrichment and chromatin accessibility at p63 ChIPseq peaks before and after YAP5SA expression in a window of −2kb to +2kb centered on the middle of the peak. D) Line plots depicting the enrichment of the H3K27Ac, H3K4me1, and H3K4me3 signal at p63 binding sites in MCF10A cells after and before YAP5SA induction.

### YAP5SA inhibits the expression of ΔNp63

We next tested whether YAP has any effect on the expression p63 that could explain the reduced chromatin-association of p63 in cells expressing YAP5SA. The induction of YAP5SA by doxycycline strongly reduced the protein expression of ΔNp63 in a time-dependent manner (Figure 7A). A robust downregulation of ΔNp63 was also observed in MCF10A cells stably expressing a hormone inducible ER-YAP2SA fusion protein (Figure 7B). After treatment of MCF10A-ER-YAP2SA with 4-OHT to activate ER-YAP2SA, levels of ΔNp63 were sharply reduced, confirming reduced protein expression of ΔNp63 by YAP. The downregulation of ΔNp63 by YAP5SA was also observed in the presence of the pharmacological proteasome inhibitor MG132, suggesting that the reduced abundance of ΔNp63 is not a consequence of its increased turnover by the proteasome (Figure 7C). As a control, MG132 stabilized p53, which is known to be regulated by ubiquitination and proteasome-dependent degradation. Because YAP5SA does not affect the protein stability of ΔNp63, we next asked whether YAP5SA regulates the mRNA levels of ΔNp63. Expression of YAP5SA significantly decreased levels of total p63 and isoform specific ΔNp63 mRNA expression while it had little effect on the mRNA expression of p53 (Figure 7D). Conversely, siRNA mediated co-depletion of endogenous YAP and the related TAZ resulted in upregulation of ΔNp63 (Figure 7E). Because YAP does not directly bind to the ΔNp63 promoter, the regulation of ΔNp63 transcription is likely an indirect effect of YAP regulating other transcription factors involved in ΔNp63 expression.

**Figure 7.**
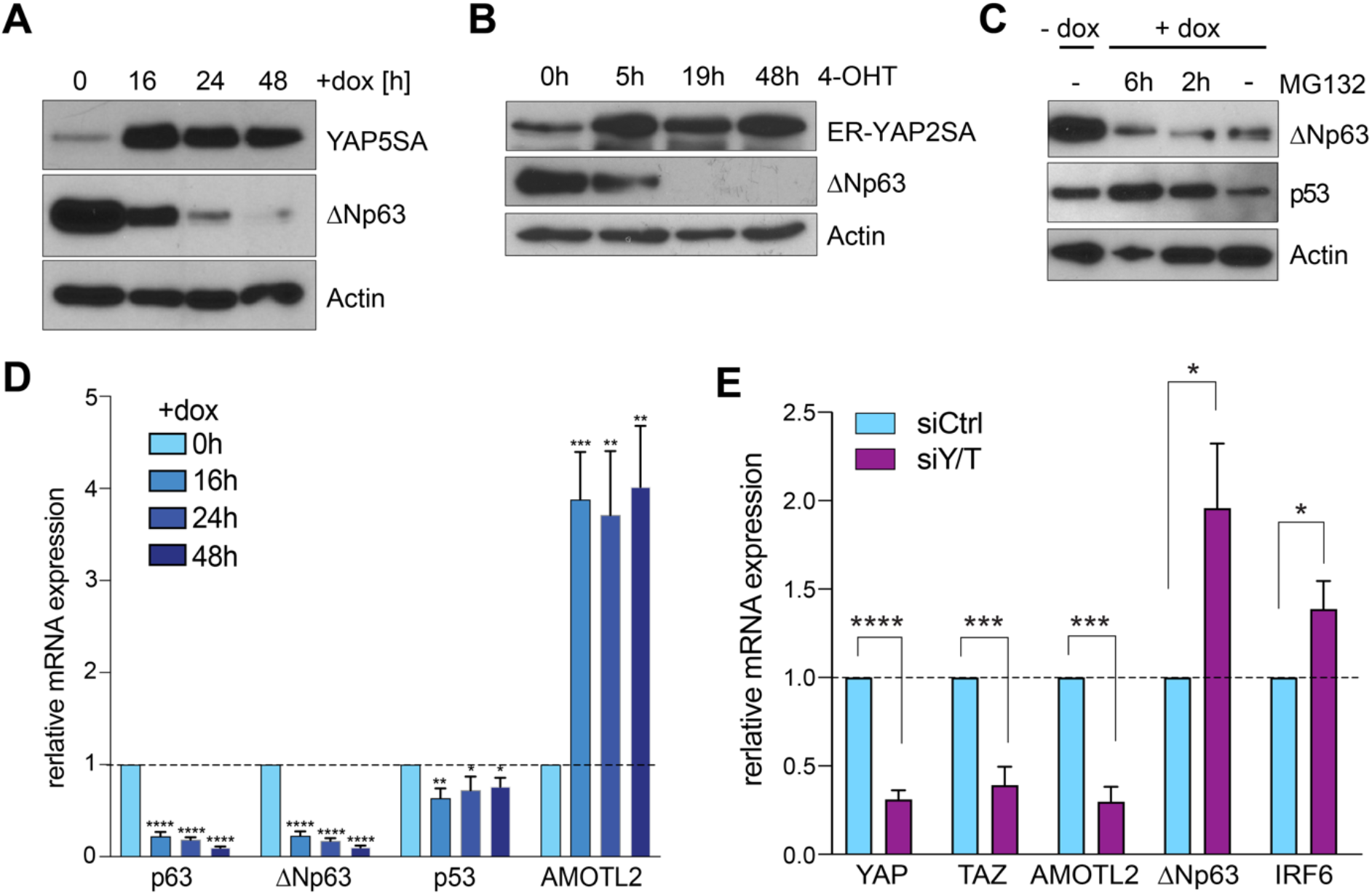
YAP5SA expression leads to downregulation of ΔNp63. A) MCF10A-YAP5SA cells were untreated (-dox) or treated with doxycycline to induce YAP5SA expression for the indicated time points. Expression of the indicated proteins was analyzed by immunoblotting. Actin served as a loading control. B). MCF10A-YAP5SA cells were untreated or treated with doxycycline to induce YAP5SA and simultaneously treated with the proteasome-inhibitor MG132. The expression of the indicated proteins was analyzed by immunoblotting. Actin was used as a loading control. C). MCF10A-ER-YAP2SA were untreated (-OHT) or treated with OHT to activate ER-YAP2SA for the indicated time points. The expression of the indicated proteins was analyzed by immunoblotting. Actin served as a loading control. D) RT-qPCR in MCF10A-YAP5SA cells before and after YAP5SA induction for the indicated time points. The expression of the indicated genes was analyzed relative to *GAPDH.* Data presented as means from biological triplicates, error bars represent SDs (n=3) E) MCF10A-YAP5SA cells were transfected with a control siRNA (siCtrl) or with siRNAs specific for YAP and TAZ (Y/T). The expression of the indicated genes was analyzed by RT-qPCR. Data presented as means from biological triplicates, error bars represent SDs (n=3). Student’s t-test. * = p<0.05, ** = p<0.01, *** = p<0.001, ns = not significant.

### Repression of ΔNp63 by YAP5SA is linked to cell migration

To find out how downregulation of p63 by YAP affects gene expression, we integrated our previous RNA-seq data of MCF10A cells expressing YAP5SA with p63 ChIP-seq data. Of the 1216 genes downregulated by YAP5SA (q<0.05), we identified 97 (8 %) genes that were also associated with a nearby p63 ChIP-seq peak. This number likely underestimates the real number of genes co-regulated by YAP and p63 as enhancers and their target genes often interact over long distances that may be missed when target gene identification is based on the nearest binding site. GO analysis showed that the 97 YAP-downregulated/ p63 bound genes were enriched for categories involved in transcription, wound healing, cell spreading, cell adhesion and cell migration (Supplemental Figure S4A).

Examples of p63-target genes that are downregulated by YAP and that are involved in cell adhesion and migration are IRF6, DLG5, MINK1 and SYNPO. Genome browser tracks of the IRF6 and MINK1 loci illustrates that YAP5SA triggers the loss of p63 binding from the enhancers, which was accompanied by reduced chromatin accessibility and reduced H3K27 acetylation (Figure 8A and Supplemental Figure S4B). ChIP-qPCR verified that p63 binding to the IRF6, DLG5, MINK1 and SYNPO and enhancers is lost upon ectopic expression of YAP5SA (Figure 8B). YAP5SA causes decreased expression of the corresponding p63-target genes while lentiviral restoration of ΔNp63 expression rescued YAP-mediated downregulation IRF6 and partially rescued MINK1 and SYNPO expression (Figure 8C). These data suggest that YAP inhibits the chromatin-binding of ΔNp63 to suppress the expression of ΔNp63 target genes. To investigate the significance of ΔNp63 downregulation for cell migration we performed transwell migration assays and found that expression of YAP5SA strongly induced migration of MCF10A cells (Figure 8D,E). Importantly, migration was rescued by restoration of ΔNp63 expression, confirming that downregulation of ΔNp63 expression is directly implicated in YAP-mediated migration. Oncogenic YAP is also known to promote mammosphere formation of MCF10A cells (9). However, and in contrast to migration, YAP-induced mammosphere formation was not rescued by ectopic ΔNp63 expression (Figure 8F,G). Altogether our data suggest that YAP inhibits ΔNp63 mRNA expression resulting in loss of ΔNp63 from enhancers, triggering a decrease in chromatin accessibility and histone acetylation at these sites and leading to the downregulation of ΔNp63 target genes. Thus, loss of ΔNp63 contributes to the global changes in the enhancer landscape upon expression of oncogenic YAP5SA. Downregulation of ΔNp63 enables cell migration in response to YAP5SA expression.

**Figure 8:**
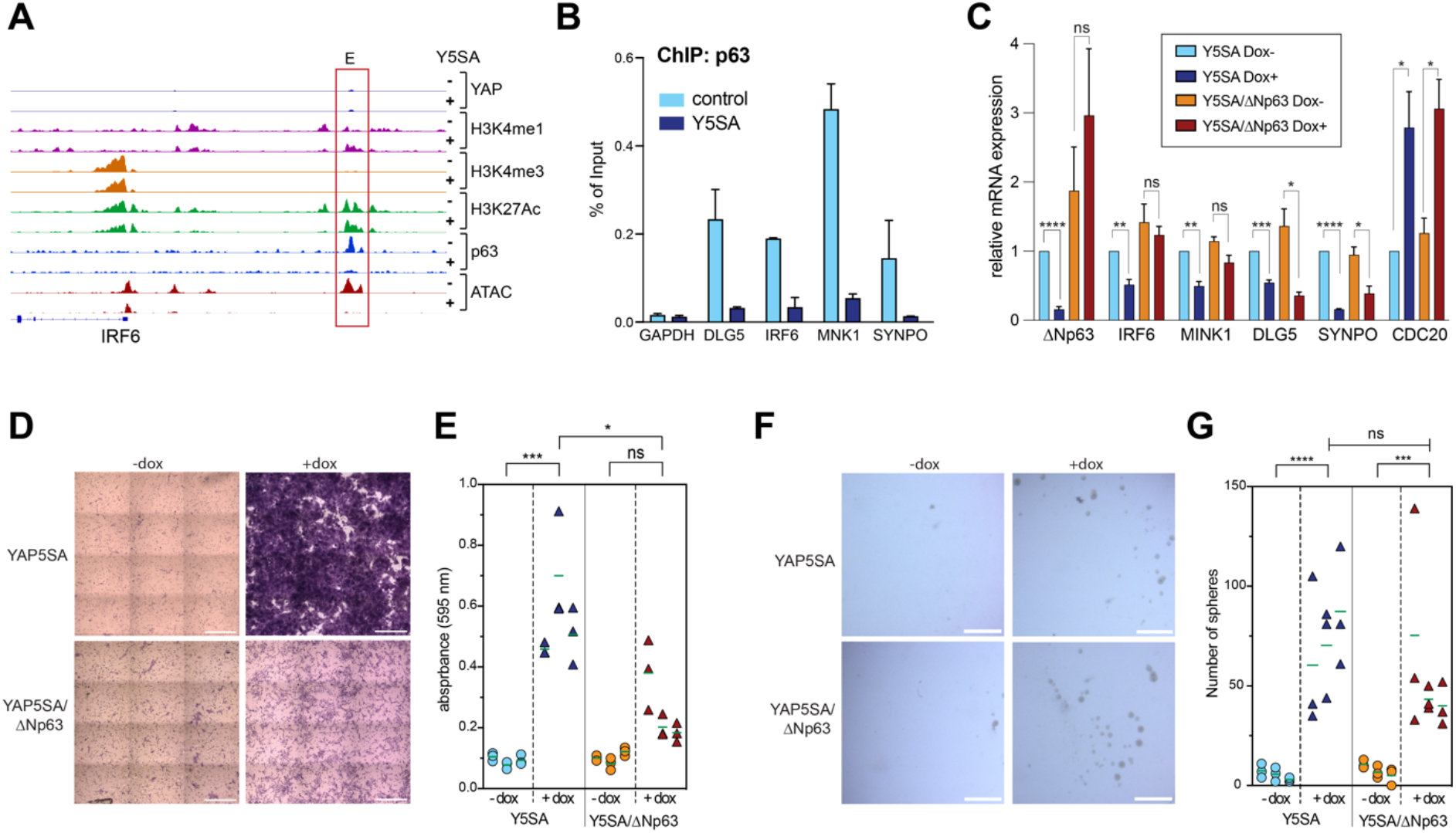
YAP5SA promotes cell migration by inhibiting ΔNp63 expression. A) Genome browser track of the *IRF6* locus, showing ChIP-seq and ATAC-seq data in control MCF10A cells (-) and after expression of YAP5SA (+). E: enhancer B) ChIP-qPCR demonstrating the binding of ΔNP63 to the enhancers of selected target genes in MCF10A-YAP5SA cells before and after the induction of YAP5SA. Mean and SDs of technical replicates of a representative experiment (n=2). C) MCF10A-YAP5SA and MCF10-YAP5SA-ΔNp63 cells were treatment with doxycycline to either induce YAP5SA or simultaneously induce YAP5SA and ΔNp63. The expression of the indicated genes was analyzed relative to *GAPDH.* Means from three independent biological replicates. Error bars represent SEM. D) Transwell migration assay of MCF10A-YAP5SA and MCF10A-YAP5SA-ΔNp63 treated as described in C. Representative images from crystal-violet stained transwell layers. Scale bar: 150 μm. E) Quantification of the transwell migration assay shown in D. Three biological replicates, each performed in technical replicates. F) Primary mammosphere formation in MCF10A-YAP5SA and MCF10-YAP5SA-ΔNp63 treated as described in C. Representative images are shown. G) Quantification of mammospheres. Mean and SDs of three biological replicates. Scale bar: 80 μm. Student’s t-test. * = p<0.05, ** = p<0.01, *** = p<0.001, ns = not significant.

## DISCUSSION

Previous studies have shown that YAP regulates gene expression by binding to distant enhancers (53). In the present study we used a well-established constitutive active allele of YAP, YAP5SA, to investigate global changes in chromatin accessibility and activity in untransformed epithelial MCF10A cells. We find that YAP5SA leads to widespread global changes in the enhancer landscape of MCF10A cells that promote the oncogenic properties of YAP. Overall YAP5SA results in thousands of newly opened and closed genome regions. These YAP-mediated chromatin changes occur relatively rapidly within two days of YAP5SA induction. By measuring the overlap of YAP binding sites with changes in accessibility and integrating these data with ChIP-seq data, we find that YAP invades a subset of partially open enhancers leading to their further opening and hyperactivation. We used the YAP target gene CDC20 to better understand the role of enhancer activation in promoting gene expression. We have previously shown that YAP binds to two enhancers stimulating the binding of the B-MYB transcription factor to the TSS of CDC20, resulting in increased expression of CDC20 (21). By CRISPRi directed at the CDC20 enhancers, we now found a link between enhancer activation by YAP and the early steps of transcription by RNA Pol II. Specifically, we demonstrate that YAP-mediated enhancer activation leads to the recruitment of RNA Pol II and the subsequent phosphorylation of Pol II at Ser5 at the TSS of CDC20. ChIP-seq confirmed that YAP increased the binding of Ser5-phosphorylated Pol II more than the Ser2 phosphorylated Pol II at MMB target genes. Phosphorylation of Pol II at Ser5 is associated with promoter escape and pausing, which have been identified as a fundamental step in transcriptional regulation (54). Our findings extend previous studies that have linked YAP to Pol II recruitment and post-recruitments steps in transcription, namely the stimulation of pause-release and productive elongation by RNA Pol II through BRD4 and CDK9 (19, 20). Transcriptional pausing puts genes in a poised state and acts as a key checkpoint that ensures the release of fully activated and mature Pol II through the promoter region allowing for rapid activation of gene expression. Pausing has also been linked to proper mRNA processing including 5’ capping and splicing of the nascent mRNA (54). We hypothesize that YAP promotes Pol II Ser5 phosphorylation in addition to controlling pause-release in order to balance initiation with elongation and to keep up with the high demand on mRNA processing due to enhanced transcription. The main kinase responsible for CTD Ser5 phosphorylation is CDK7, a subunit of the general transcription factor TFIIH. While co-immunoprecipitations did not show a direct biochemical interaction between CDK7 and MMB, ChIP and PLA experiments provide evidence for increased YAP-dependent recruitment of CDK7 to MMB target genes. Chemical inhibition of CDK7 abolished the hyperinduction of MMB-target genes, although it is important to consider that CDK7 inhibition may have global effects on gene transcription. Although CDK7 has recently been shown to phosphorylate YAP in the nucleus and to prevent its proteasomal degradation (50), in our experimental system, we did not observe any effect of CDK7 inhibition on YAP expression. It is therefore unlikely that the dependence of MMB-target gene expression by YAP on CDK7 is a consequence of the previously described role of CDK7 in YAP turnover. Activation of CDC20 was not only associated with increased RNA Pol II phosphorylation, but was also accompanied by histone acetylation at the promoter of this gene. Since H3K9 acetylation at the CDC20 promoter was dependent on enhancer activation, GCN5/PCAF-containing SAGA and ATAC complexes may have a more direct role in MMB target gene activation than other histone acetyltransferases as they are known to catalyze this modification (55).

In addition to enhancer activation, we also identified regions with decreased chromatin accessibility in YAP5SA expressing cells, which is explained, at least in part, by a previously unknown function of YAP in downregulating ΔNp63. p63, which plays an important role in mammary epithelial development and self-renewal can be expressed as two isoforms, TAp63 and ΔNp63 (56, 57). While ΔNp63 functions as an oncogene by inhibiting the function of p53, TAp63 and TAp73, there is also evidence that reduced expression of ΔNp63α plays roles in EMT, cell motility and cancer metastasis (58–63). ΔNp63 is an unstable protein that is rapidly turned over by proteasomal degradation (64). Although it has previously been reported that YAP physically interacts with ΔNp63 in JHU-22 cells to reduce its half-life (65), YAP does not regulate ΔNp63 protein stability in MCF10A cells, but inhibits the transcription of the ΔNp63 mRNA. The inhibition of ΔNp63 expression by YAP is reminiscent of down-regulation of ΔNp63 expression by oncogenic H-Ras, PI3-K and HER2 signaling (58, 59). As it has previously been reported that YAP can suppress PTEN (66), it is tempting to speculate that PI3-K pathway activation contributes to ΔNp63 suppression by YAP. Notably, ΔNp63 binds to its own intronic enhancer to maintain its sustained expression (67). Thus, the signals that lead to downregulation of ΔNp63 expression could be transient and once the positive feedback loop has been interrupted, ΔNp63 could be permanently silenced. More work will be required to determine how YAP suppresses ΔNp63 expression. More importantly, we show that repression of ΔNp63 is pivotal for YAP-induced cell migration. By integrating ChIP-seq and RNA-seq data we identified a group of enhancer-associated genes regulated by ΔNp63 and YAP5SA, many of which have previously been shown to be involved in cell migration and adhesion. One example is interferon regulatory factor 6 (IRF6) which was strongly downregulated by YAP5SA and which has been shown to inhibit migration and cell invasion in squamous cell carcinomas and colorectal cancer cells (68, 69). Further investigation is required to determine whether suppression of IRF6 contributes to YAP-mediated migration.

In conclusion, we have analyzed global changes in the chromatin status by oncogenic YAP in untransformed MCF10A cells and linked these changes to the expression of genes relevant for cell cycle regulation and migration (Figure 9). Altogether, our findings point to oncogenic activities driven by YAP that may have relevance in cancer biology.

**Figure 9:**
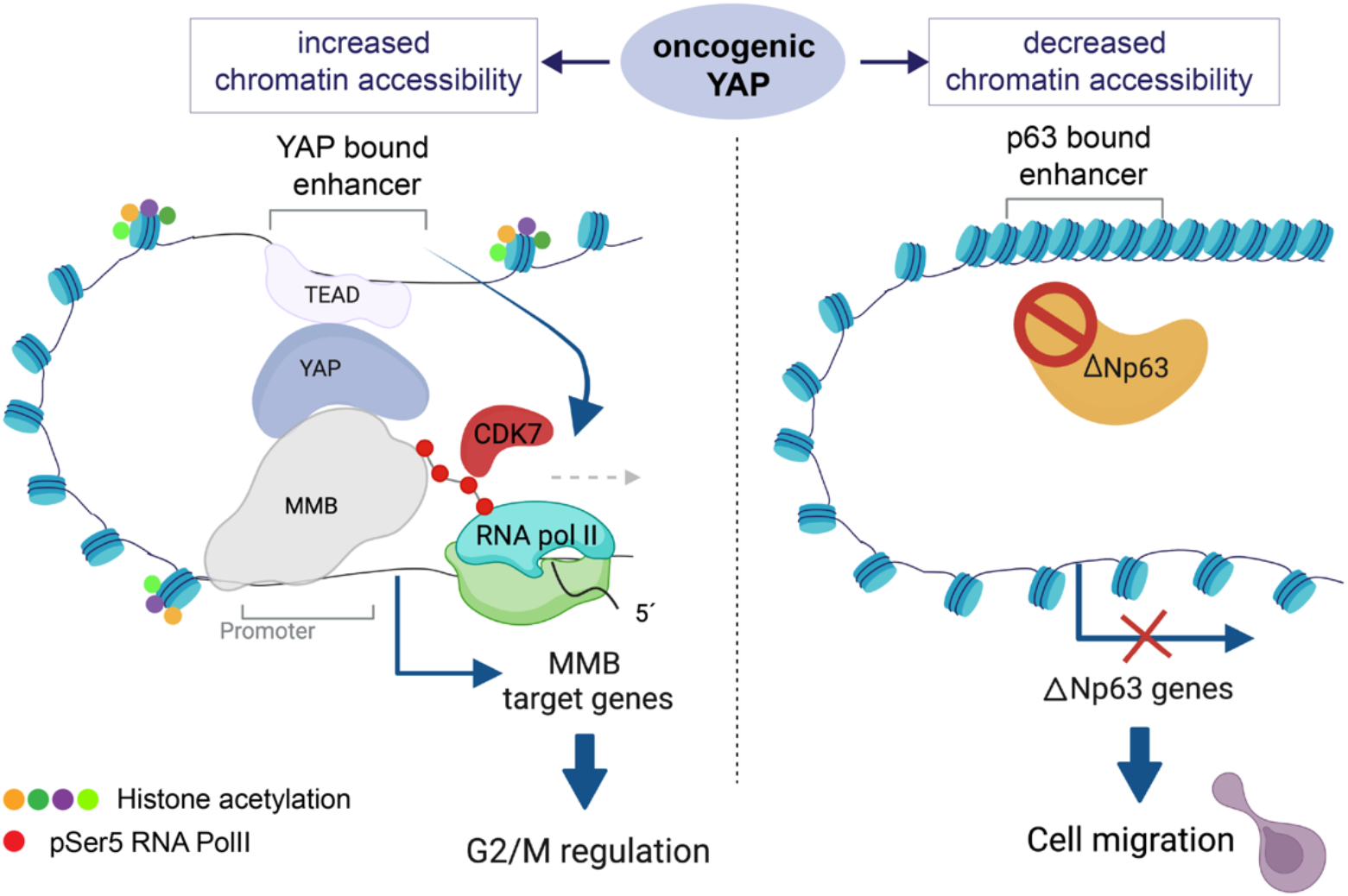
Summary of the results and model. Oncogenic YAP5SA invades a subset of partially accessible enhancers leading to their further opening and hyperactivation. YAP-mediated enhancer activation promotes the recruitment of RNA Pol II and the subsequent CDK7-dependent phosphorylation of Pol II at Ser5 at the TSS of MMB-regulated promoters of cell cycle genes. YAP5SA also leads to less accessible “closed” chromatin regions, which are not directly YAP-bound but which contain binding motifs for p63. Diminished accessibility at these regions is a consequence of reduced expression and chromatin-binding of ΔNp63 resulting in downregulation of ΔNp63-target genes and promoting YAP-mediated cell migration.

## DATA AVAILIBILITY

ATAC-seq and ChIP-sequencing datasets are available at the NCBI’s Gene Expression Omnibus (70) under the accession number: GSE193704.

## ACKNOWLEDGEMENTS

We thank Svenja Meierjohann for sharing reagents. We thank Anna-Lena Mattes and Susi Spahr for assistance with experiments. Some figure panels were created with Biorender.

## AUTHOR CONTRIBUTION

SG planned the study; SG, MCF, DG, and FL designed experiments; SG, MCF, DG, FL, JT, BG, GW and NS conducted the experiments; SG, MCF, DG, JT and FL analyzed data; SG and SK performed bioinformatic analyses; CPA performed and supervised next generation sequencing; SG and MCF wrote the manuscript.

## REFERENCES

1. Totaro,A., Panciera,T. and Piccolo,S. (2018) YAP/TAZ upstream signals and downstream responses. Nature cell biology, 20, 888–899.

2. Moya,I.M. and Halder,G. (2019) Hippo–YAP/TAZ signalling in organ regeneration and regenerative medicine. Nat Rev Mol Cell Bio, 20, 211–226.

3. Zanconato,F., Battilana,G., Cordenonsi,M. and Piccolo,S. (2016) YAP/TAZ as therapeutic targets in cancer. Current opinion in pharmacology, 29, 26–33.

4. Yu,F.-X., Zhao,B. and Guan,K.-L. (2015) Hippo Pathway in Organ Size Control, Tissue Homeostasis, and Cancer. Cell, 163, 811–828.

5. Meng,Z., Moroishi,T. and Guan,K.-L. (2016) Mechanisms of Hippo pathway regulation. Genes & development, 30, 1–17.

6. Piccolo,S., Dupont,S. and Cordenonsi,M. (2014) The biology of YAP/TAZ: hippo signaling and beyond. Physiological reviews, 94, 1287–1312.

7. Thompson,B.J. (2020) YAP/TAZ: Drivers of Tumor Growth, Metastasis, and Resistance to Therapy. Bioessays, 42, 1900162.

8. Stein,C., Bardet,A.F., Roma,G., Bergling,S., Clay,I., Ruchti,A., Agarinis,C., Schmelzle,T., Bouwmeester,T., Schübeler,D., et al. (2015) YAP1 Exerts Its Transcriptional Control via TEAD-Mediated Activation of Enhancers. PLoS genetics, 11, e1005465.

9. Zanconato,F., Forcato,M., Battilana,G., Azzolin,L., Quaranta,E., Bodega,B., Rosato,A., Bicciato,S., Cordenonsi,M. and Piccolo,S. (2015) Genome-wide association between YAP/TAZ/TEAD and AP-1 at enhancers drives oncogenic growth. Nature cell biology, 17, 1218–1227.

10. Zhu,C., Li,L., Zhang,Z., Bi,M., Wang,H., Su,W., Hernandez,K., Liu,P., Chen,J., Chen,M., et al. (2019) A Non-canonical Role of YAP/TEAD Is Required for Activation of Estrogen-Regulated Enhancers in Breast Cancer. Mol Cell, 75, 791–806.e8.

11. Cebola,I., Rodríguez-Seguí,S.A., Cho,C.H.-H., Bessa,J., Rovira,M., Luengo,M., Chhatriwala,M., Berry,A., Ponsa-Cobas,J., Maestro,M.A., et al. (2015) TEAD and YAP regulate the enhancer network of human embryonic pancreatic progenitors. Nature cell biology, 17, 615–626.

12. Bulger,M. and Groudine,M. (2010) Enhancers: The abundance and function of regulatory sequences beyond promoters. Dev Biol, 339, 250–257.

13. Hnisz,D., Abraham,B.J., Lee,T.I., Lau,A., Saint-André,V., Sigova,A.A., Hoke,H.A. and Young,R.A. (2013) Super-Enhancers in the Control of Cell Identity and Disease. Cell, 155, 934–947.

14. Roe,J.-S., Hwang,C.-I., Somerville,T.D.D., Milazzo,J.P., Lee,E.J., Silva,B.D., Maiorino,L., Tiriac,H., Young,C.M., Miyabayashi,K., et al. (2017) Enhancer Reprogramming Promotes Pancreatic Cancer Metastasis. Cell, 10.1016/j.cell.2017.07.007.

15. Kron,K.J., Bailey,S.D. and Lupien,M. (2014) Enhancer alterations in cancer: a source for a cell identity crisis. Genome Med, 6, 77.

16. Bradner,J.E., Hnisz,D. and Young,R.A. (2017) Transcriptional Addiction in Cancer. Cell, 168, 629–643.

17. Liu,X., Li,H., Rajurkar,M., Li,Q., Cotton,J.L., Ou,J., Zhu,L.J., Goel,H.L., Mercurio,A.M., Park,J.-S., et al. (2016) Tead and AP1 Coordinate Transcription and Motility. Cell reports, 14, 1169–1180.

18. He,L., Pratt,H., Gao,M., Wei,F., Weng,Z. and Struhl,K. (2021) YAP and TAZ are transcriptional co-activators of AP-1 proteins and STAT3 during breast cellular transformation. Elife, 10, e67312.

19. Galli,G.G., Carrara,M., Yuan,W.-C., Valdes-Quezada,C., Gurung,B., Pepe-Mooney,B., Zhang,T., Geeven,G., Gray,N.S., Laat,W. de, et al. (2015) YAP Drives Growth by Controlling Transcriptional Pause Release from Dynamic Enhancers. Molecular cell, 60, 328–337.

20. Zanconato,F., Battilana,G., Forcato,M., Filippi,L., Azzolin,L., Manfrin,A., Quaranta,E., Biagio,D.D., Sigismondo,G., Guzzardo,V., et al. (2018) Transcriptional addiction in cancer cells is mediated by YAP/TAZ through BRD4. Nature medicine, 24, 1599–1610.

21. Pattschull,G., Walz,S., Gründl,M., Schwab,M., Rühl,E., Baluapuri,A., Cindric-Vranesic,A., Kneitz,S., Wolf,E., Ade,C.P., et al. (2019) The Myb-MuvB Complex Is Required for YAP-Dependent Transcription of Mitotic Genes. Cell Reports, 27, 3533–3546.e7.

22. Gründl,M., Walz,S., Hauf,L., Schwab,M., Werner,K.M., Spahr,S., Schulte,C., Maric,H.M., Ade,C.P. and Gaubatz,S. (2020) Interaction of YAP with the Myb-MuvB (MMB) complex defines a transcriptional program to promote the proliferation of cardiomyocytes. Plos Genet, 16, e1008818.

23. Sadasivam,S., Duan,S. and DeCaprio,J.A. (2012) The MuvB complex sequentially recruits B-Myb and FoxM1 to promote mitotic gene expression. Genes & development, 26, 474–489.

24. Pilkinton,M., Sandoval,R., Song,J., Ness,S.A. and Colamonici,O.R. (2007) Mip/LIN-9 regulates the expression of B-Myb and the induction of cyclin A, cyclin B, and CDK1. The Journal of biological chemistry, 282, 168–175.

25. Pilkinton,M., Sandoval,R. and Colamonici,O.R. (2007) Mammalian Mip/LIN-9 interacts with either the p107, p130/E2F4 repressor complex or B-Myb in a cell cycle-phase-dependent context distinct from the Drosophila dREAM complex. Oncogene, 26, 7535–7543.

26. Schmit,F., Korenjak,M., Mannefeld,M., Schmitt,K., Franke,C., Eyss,B. von, Gagrica,S., Hänel,F., Brehm,A. and Gaubatz,S. (2007) LINC, a human complex that is related to pRB-containing complexes in invertebrates regulates the expression of G2/M genes. Cell cycle (Georgetown, Tex), 6, 1903–1913.

27. Osterloh,L., Eyss,B. von, Schmit,F., Rein,L., Hübner,D., Samans,B., Hauser,S. and Gaubatz,S. (2007) The human synMuv-like protein LIN-9 is required for transcription of G2/M genes and for entry into mitosis. The EMBO journal, 26, 144–157.

28. Wei,T., Weiler,S.M.E., Tóth,M., Sticht,C., Lutz,T., Thomann,S., Torre,C.D.L., Straub,B., Merker,S., Ruppert,T., et al. (2019) YAP-dependent induction of UHMK1 supports nuclear enrichment of the oncogene MYBL2 and proliferation in liver cancer cells. Oncogene,10.1038/s41388-019-0801-y.

29. Eyss,B. von, Jaenicke,L.A., Kortlever,R.M., Royla,N., Wiese,K.E., Letschert,S., McDuffus,L.-A., Sauer,M., Rosenwald,A., Evan,G.I., et al. (2015) A MYC-Driven Change in Mitochondrial Dynamics Limits YAP/TAZ Function in Mammary Epithelial Cells and Breast Cancer. Cancer cell, 28, 743–757.

30. Buenrostro,J.D., Giresi,P.G., Zaba,L.C., Chang,H.Y. and Greenleaf,W.J. (2013) Transposition of native chromatin for fast and sensitive epigenomic profiling of open chromatin, DNA-binding proteins and nucleosome position. Nature methods, 10, 1213–1218.

31. Skene,P.J. and Henikoff,S. (2017) An efficient targeted nuclease strategy for high-resolution mapping of DNA binding sites. Elife, 6, e21856.

32. Meers,M.P., Bryson,T.D., Henikoff,J.G. and Henikoff,S. (2019) Improved CUTRUN chromatin profiling tools. Elife, 8, e46314.

33. Liu,N., Hargreaves,V.V., Zhu,Q., Kurland,J.V., Hong,J., Kim,W., Sher,F., Macias-Trevino,C., Rogers,J.M., Kurita,R., et al. (2018) Direct Promoter Repression by BCL11A Controls the Fetal to Adult Hemoglobin Switch. Cell, 173, 430–442.e17.

34. Kearns,N.A., Genga,R.M.J., Enuameh,M.S., Garber,M., Wolfe,S.A. and Maehr,R. (2013) Cas9 effector-mediated regulation of transcription and differentiation in human pluripotent stem cells. Development, 141, 219–223.

35. Doench,J.G., Fusi,N., Sullender,M., Hegde,M., Vaimberg,E.W., Donovan,K.F., Smith,I., Tothova,Z., Wilen,C., Orchard,R., et al. (2016) Optimized sgRNA design to maximize activity and minimize off-target effects of CRISPR-Cas9. Nature biotechnology, 34, 184–191.

36. Stringer,B.W., Day,B.W., D’Souza,R.C.J., Jamieson,P.R., Ensbey,K.S., Bruce,Z.C., Lim,Y.C., Goasdoué,K., Offenhäuser,C., Akgül,S., et al. (2019) A reference collection of patient-derived cell line and xenograft models of proneural, classical and mesenchymal glioblastoma. Sci Rep-uk, 9, 4902.

37. Afgan,E., Baker,D., Batut,B., Beek,M. van den, Bouvier,D., Čech,M., Chilton,J., Clements,D., Coraor,N., Grüning,B.A., et al. (2018) The Galaxy platform for accessible, reproducible and collaborative biomedical analyses: 2018 update. Nucleic Acids Res, 46, W537–W544.

38. Langmead,B. and Salzberg,S.L. (2012) Fast gapped-read alignment with Bowtie 2. Nature methods, 9, 357–359.

39. Liu,R., Holik,A.Z., Su,S., Jansz,N., Chen,K., Leong,H.S., Blewitt,M.E., Asselin-Labat,M.-L., Smyth,G.K. and Ritchie,M.E. (2015) Why weight? Modelling sample and observational level variability improves power in RNA-seq analyses. Nucleic acids research, 43, e97–e97.

40. Schep,A.N., Buenrostro,J.D., Denny,S.K., Schwartz,K., Sherlock,G. and Greenleaf,W.J. (2015) Structured nucleosome fingerprints enable high-resolution mapping of chromatin architecture within regulatory regions. Genome Res, 25, 1757–1770.

41. Ramírez,F., Ryan,D.P., Grüning,B., Bhardwaj,V., Kilpert,F., Richter,A.S., Heyne,S., Dündar,F. and Manke,T. (2016) deepTools2: a next generation web server for deep-sequencing data analysis. Nucleic Acids Res, 44, W160–5.

42. Fischer,M., Quaas,M., Steiner,L. and Engeland,K. (2016) The p53-p21-DREAM-CDE/CHR pathway regulates G2/M cell cycle genes. Nucleic acids research, 44, 164–174.

43. Robinson,J.T., Thorvaldsdóttir,H., Winckler,W., Guttman,M., Lander,E.S., Getz,G. and Mesirov,J.P. (2011) Integrative genomics viewer. Nature biotechnology, 29, 24–26.

44. Schep,A.N., Wu,B., Buenrostro,J.D. and Greenleaf,W.J. (2017) chromVAR: inferring transcription-factor-associated accessibility from single-cell epigenomic data. Nat Methods, 14, 975–978.

45. Glover-Cutter,K., Larochelle,S., Erickson,B., Zhang,C., Shokat,K., Fisher,R.P. and Bentley,D.L. (2009) TFIIH-Associated Cdk7 Kinase Functions in Phosphorylation of C-Terminal Domain Ser7 Residues, Promoter-Proximal Pausing, and Termination by RNA Polymerase II ^▿^†. Mol Cell Biol, 29, 5455–5464.

46. Trizzino,M., Barbieri,E., Petracovici,A., Wu,S., Welsh,S.A., Owens,T.A., Licciulli,S., Zhang,R. and Gardini,A. (2018) The Tumor Suppressor ARID1A Controls Global Transcription via Pausing of RNA Polymerase II. Cell Reports, 23, 3933–3945.

47. Branigan,T.B., Kozono,D., Schade,A.E., Deraska,P., Rivas,H.G., Sambel,L., Reavis,H.D., Shapiro,G.I., D’Andrea,A.D. and DeCaprio,J.A. (2021) MMB-FOXM1-driven premature mitosis is required for CHK1 inhibitor sensitivity. Cell Reports, 34, 108808.

48. Saldivar,J.C., Hamperl,S., Bocek,M.J., Chung,M., Bass,T.E., Cisneros-Soberanis,F., Samejima,K., Xie,L., Paulson,J.R., Earnshaw,W.C., et al. (2018) An intrinsic S/G2 checkpoint enforced by ATR. Science, 361, 806–810.

49. Brodowska,K., Al-Moujahed,A., Marmalidou,A., Horste,M.M.Z., Cichy,J., Miller,J.W., Gragoudas,E. and Vavvas,D.G. (2014) The clinically used photosensitizer Verteporfin (VP) inhibits YAP-TEAD and human retinoblastoma cell growth in vitro without light activation. Experimental eye research, 124, 67–73.

50. Cho,Y.S., Li,S., Wang,X., Zhu,J., Zhuo,S., Han,Y., Yue,T., Yang,Y. and Jiang,J. (2020) CDK7 regulates organ size and tumor growth by safeguarding the Hippo pathway effector Yki/Yap/Taz in the nucleus. Gene Dev, 34, 53–71.

51. Carroll,D.K., Carroll,J.S., Leong,C.-O., Cheng,F., Brown,M., Mills,Alea.A., Brugge,J.S. and Ellisen,L.W. (2006) p63 regulates an adhesion programme and cell survival in epithelial cells. Nat Cell Biol, 8, 551–561.

52. Uzunbas,G.K., Ahmed,F. and Sammons,M.A. (2019) Control of p53-dependent transcription and enhancer activity by the p53 family member p63. J Biol Chem, 294, 10720–10736.

53. Lopez-Hernandez,A., Sberna,S. and Campaner,S. (2021) Emerging Principles in the Transcriptional Control by YAP and TAZ. Cancers, 13, 4242.

54. Core,L. and Adelman,K. (2019) Promoter-proximal pausing of RNA polymerase II: a nexus of gene regulation. Gene Dev, 33, 960–982.

55. Helmlinger,D. and Tora,L. (2017) Sharing the SAGA. Trends Biochem Sci, 42, 850–861.

56. Su,X., Chakravarti,D. and Flores,E.R. (2013) p63 steps into the limelight: crucial roles in the suppression of tumorigenesis and metastasis. Nat Rev Cancer, 13, 136–143.

57. Bergholz,J. and Xiao,Z.-X. (2012) Role of p63 in Development, Tumorigenesis and Cancer Progression. Cancer Microenviron, 5, 311–322.

58. Yoh,K.E., Regunath,K., Guzman,A., Lee,S.-M., Pfister,N.T., Akanni,O., Kaufman,L.J., Prives,C. and Prywes,R. (2016) Repression of p63 and induction of EMT by mutant Ras in mammary epithelial cells. Proc National Acad Sci, 113, E6107–E6116.

59. Hu,L., Liang,S., Chen,H., Lv,T., Wu,J., Chen,D., Wu,M., Sun,S., Zhang,H., You,H., et al. (2017) ΔNp63α is a common inhibitory target in oncogenic PI3K/Ras/Her2-induced cell motility and tumor metastasis. Proc National Acad Sci, 114, E3964–E3973.

60. Barbieri,C.E., Tang,L.J., Brown,K.A. and Pietenpol,J.A. (2006) Loss of p63 Leads to Increased Cell Migration and Up-regulation of Genes Involved in Invasion and Metastasis. Cancer Res, 66, 7589–7597.

61. Tucci,P., Agostini,M., Grespi,F., Markert,E.K., Terrinoni,A., Vousden,K.H., Muller,P.A.J., Dötsch,V., Kehrloesser,S., Sayan,B.S., et al. (2012) Loss of p63 and its microRNA-205 target results in enhanced cell migration and metastasis in prostate cancer. Proc National Acad Sci, 109, 15312–15317.

62. Zhao,W., Wang,H., Han,X., Ma,J., Zhou,Y., Chen,Z., Zhou,H., Xu,H., Sun,Z., Kong,B., et al. (2016) ΔNp63α attenuates tumor aggressiveness by suppressing miR-205/ZEB1-mediated epithelial–mesenchymal transition in cervical squamous cell carcinoma. Tumor Biol, 37, 10621–10632.

63. Lindsay,J., McDade,S.S., Pickard,A., McCloskey,K.D. and McCance,D.J. (2010) Role of DeltaNp63gamma in epithelial to mesenchymal transition. J Biological Chem, 286, 3915–24.

64. Galli,F., Rossi,M., D’Alessandra,Y., Simone,M.D., Lopardo,T., Haupt,Y., Alsheich-Bartok,O., Anzi,S., Shaulian,E., Calabrò,V., et al. (2010) MDM2 and Fbw7 cooperate to induce p63 protein degradation following DNA damage and cell differentiation. J Cell Sci, 123, 2423–2433.

65. Chatterjee,A., Sen,T., Chang,X. and Sidransky,D. (2010) Yes-associated protein 1 regulates the stability of ΔNp63α. Cell Cycle, 9, 162–167.

66. Tumaneng,K., Schlegelmilch,K., Russell,R.C., Yimlamai,D., Basnet,H., Mahadevan,N., Fitamant,J., Bardeesy,N., Camargo,F.D. and Guan,K.-L. (2012) YAP mediates crosstalk between the Hippo and PI(3)K–TOR pathways by suppressing PTEN via miR-29. Nat Cell Biol, 14, 1322–1329.

67. Antonini,D., Rossi,B., Han,R., Minichiello,A., Palma,T.D., Corrado,M., Banfi,S., Zannini,M., Brissette,J.L. and Missero,C. (2006) An Autoregulatory Loop Directs the Tissue-Specific Expression of p63 through a Long-Range Evolutionarily Conserved Enhancer†. Mol Cell Biol, 26, 3308–3318.

68. Tan,L., Qu,W., Wu,D., Liu,M., Ai,Q., Hu,H., Wang,Q., Chen,W. and Zhou,H. (2022) The interferon regulatory factor 6 promotes cisplatin sensitivity in colorectal cancer. Bioengineered, 13, 10504–10517.

69. Botti,E., Spallone,G., Moretti,F., Marinari,B., Pinetti,V., Galanti,S., Meo,P.D.D., Nicola,F.D., Ganci,F., Castrignanò,T., et al. (2011) Developmental factor IRF6 exhibits tumor suppressor activity in squamous cell carcinomas. Proc National Acad Sci, 108, 13710–13715.

70. Edgar,R., Domrachev,M. and Lash,A.E. (2002) Gene Expression Omnibus: NCBI gene expression and hybridization array data repository. Nucleic acids research, 30, 207–210.

