## Supplemental Material for "Oncogenic YAP mediates changes in chromatin accessibility and activity that drive cell cycle gene expression and cell migration"

#### SUPPLEMENTAL FIGURE AND TABLE LEGENDS

**Supplemental Figure S1:** A) Line plots depicting chromatin accessibility (ATAC), nucleosome signal (NucleoATAC), and the levels of the indicated proteins by ChIPseq at the TSS of YAP-MMB target genes in MCF10A cells with and without expression of YAP5SA. B) Line plots depicting chromatin accessibility (ATAC), nucleosome signal (NucleoATAC), and the levels of the indicated proteins by ChIPseq at the TSS of genes with gained ATAC-peaks in their promoter with and without expression of YAP5SA. C) Genome browser tracks showing chromatin accessibility and YAP-binding to two *CDC20* enhancers (E1 and E2). Long-range interactions between the *CDC20* promoter and enhancers as determined by 4C-seq are indicated by arrows (Pattschull et al., 2019). D) Gene browser tracks of the *CDC20* TSS and enhancer regions depicting the indicating ChIPseq and ATACseq data before and after expression of YAP5SA.

**Supplemental Figure S2:** A) Expression of Cas9-KRAB and YAP5SA in MCF10A cells before and after induction with doxycycline was analyzed by immunoblotting. Actin served as a control. B) Expression of Cas9-KRAB was analyzed by immunostaining with an HA-antibody. Nuclei were stained with Hoechst. +dox: cells were treated with doxycycline to induce the expression of Cas9-KRAB and YAP5SA

**Supplemental Figure S3:** A) Immunoblotting of the indicating proteins in lysates of cells treated as described in Figure 5B demonstrating that CDK7i does not affect the levels of YAP. B) Nuclear lysates of MCF10A-YAP5SA cells treated with or without doxycycline were subjected to immunoprecipitation with CDK7 antibodies. Bound proteins were detected by immunoblotting. XPD was used as positive control. Immunoprecipitations with nonspecific IgG served as a negative control.

**Supplemental Figure S4:** A) GO analysis of YAP and p63 co-regulated genes that were identified by integrating p63 ChIP-seq and YAP5SA RNA-seq data sets. The identified genes are involved in diverse biological processes including development and wound healing. B) Genome browser track of the *MINK1* locus, showing ChIP-seq and ATAC-seq data in control MCF10A cells (-) and after expression of YAP5SA (+). E: enhancer

**Supplemental Table 1:** List of oligonucleotide and siRNA sequences

**Supplemental Table 2:** List of antibodies used in this study

Supplemental Figure S1

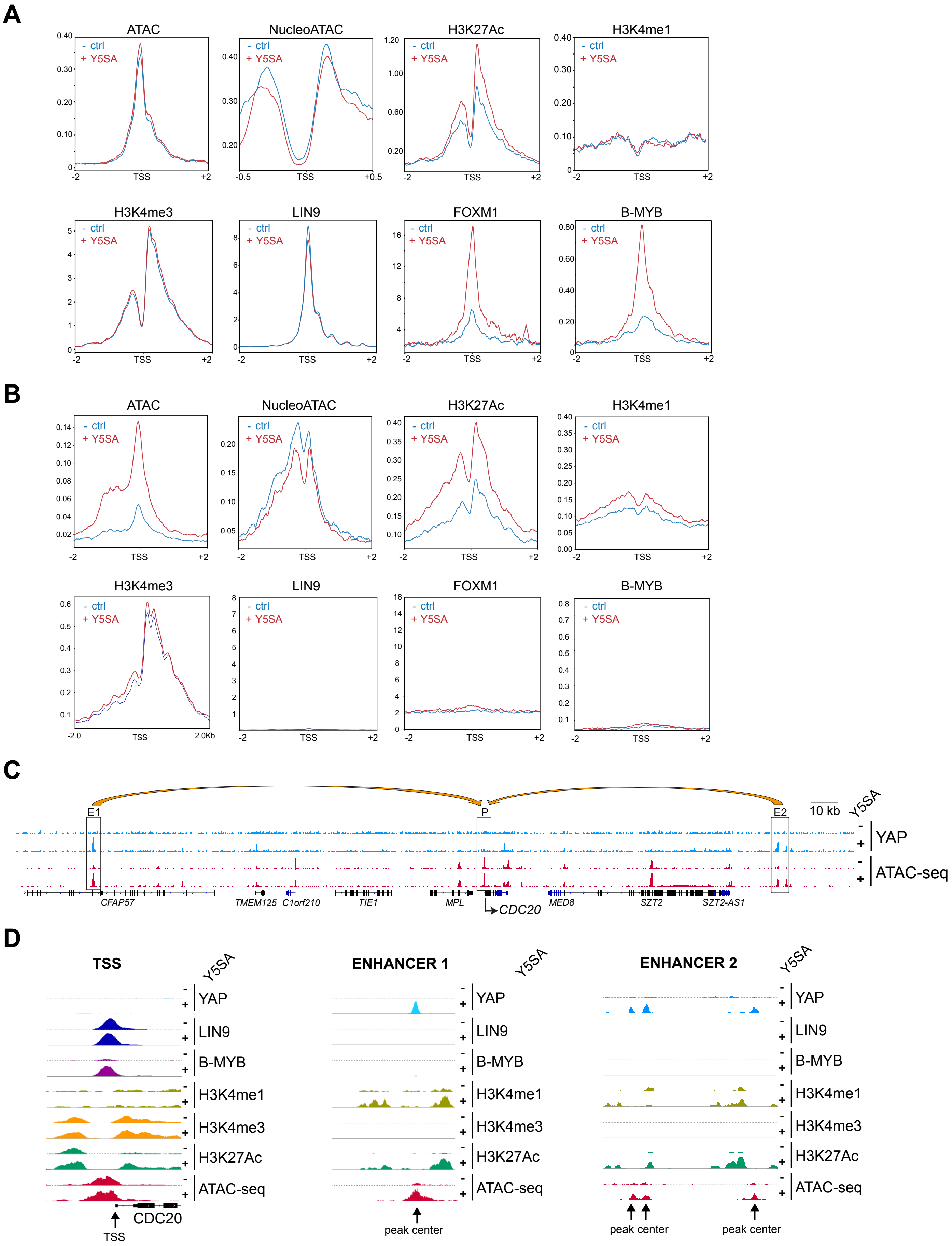

### Supplemental Figure S2

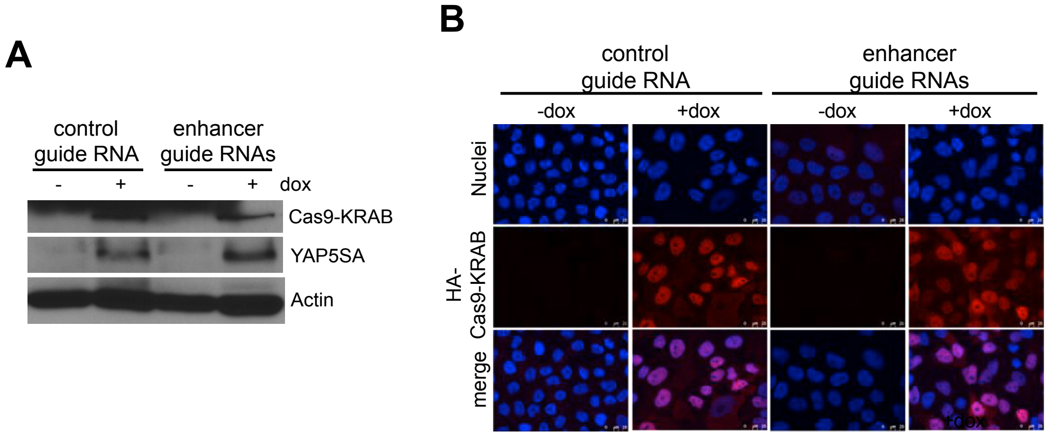

### Supplemental Figure S3

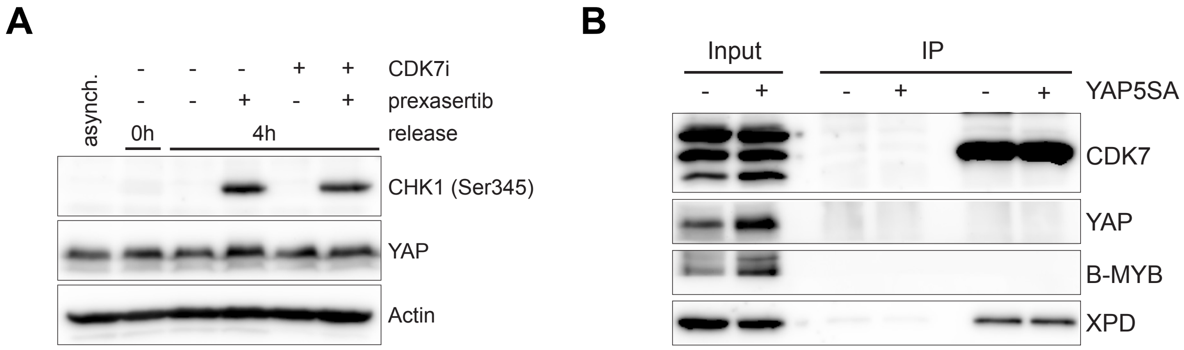

### Supplemental Figure S4

A

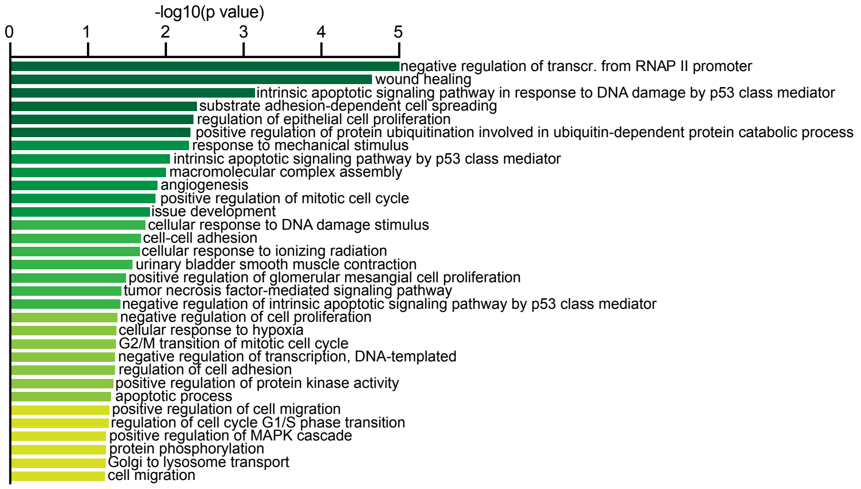


B

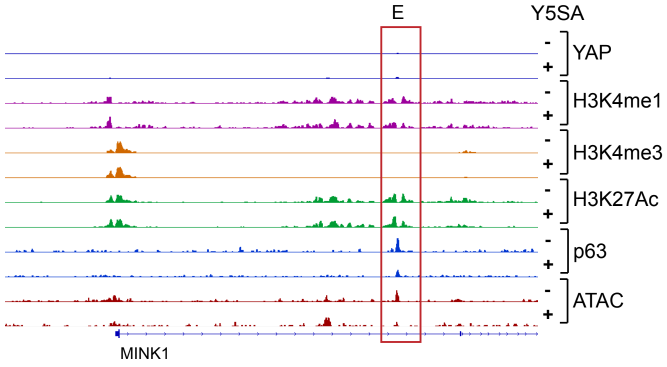

**Supplemental Table 1: Oligonucleotide and siRNA sequences**

**ATAC-seq**

| Name | Sequence |
|---|---|
| Ad1_noMX | AATGATACGGCGACCACCGAGATCTACACTCGTCGGCAGCGTCAGATGT*G |
| Ad2.1 | CAAGCAGAAGACGGCATACGAGATTCGCCTTAGTCTCGTGGGCTCGGAGATG*T |
| Ad2.2 | CAAGCAGAAGACGGCATACGAGATCTAGTACGGTCTCGTGGGCTCGGAGATG*T |
| Ad2.3 | CAAGCAGAAGACGGCATACGAGATTTCTGCCTGTCTCGTGGGCTCGGAGATG*T |
| Ad2.4 | CAAGCAGAAGACGGCATACGAGATGCTCAGGAGTCTCGTGGGCTCGGAGATG*T |

**ChIP-qPCR**

| Region | fw primer | bw primer |
|---|---|---|
| CDC20 -428 | GGCTCTCCTTCCCCTTCTAG | TCCGAGAACCTGCAGAAGTT |
| CDC20 -41 | GGTTGCGACGGTTGGATTTT | CTTTAACACGCCTGGCTTACG |
| CDC20 +120 | GGTGGCCCTGATTTTGTGG | CTGGAGGAAAGGAGGCGAC |
| CDC20 +1827 | TCCACCACCATGATGTTCGG | CACTGGCCAAATGTCGTCCA |
| CDC20 +4020 | GAGTCTGACCATGAGCCCAG | CTCAGGGTCTCATCTGCTGC |
| CDC20 +4113 | ATGGCGCTGTTTTGAGTTGG | ACTGAGGTGATGGGTTGGTC |
| CDC20 TES | ACCTTTGGCCCAGAAGCTAC | CAAAGGGTCTCGTGGCCTT |
| CDC20 enhancer 1 | CCTGCAGAGTGAGCCATGTG | GAGCTGGCTGTGTCCTTTGA |
| CDC20 enhancer 2 | AAATACTCTTGGCCACGCAC | AGTCTCACCATGCTGCCTAG |
| DLG5 enhancer | TGAGTGCTCCTGAGGAAGTG | TCTACTTCCTTACCGCCTGC |
| IRF6 enhancer | TCCTTGACCCTGATTGAGCC | TCCAGACATTGACGCAGGTA |
| MINK1 enhancer | TCACACATCCCTCTGGCATA | AGAAGGAAGGCGAAGAAACC |
| SYNPO enhancer | CACGAGTCCCGTGTACCTG | AGCCTGTGCTACACACAGCA |

**RT-qPCR primers**

| Gene | fw primer | bw primer |
|---|---|---|
| AMOTL2 | AGGCTGCAGAGAGACAATGAG | CTCAGAGAGCCGCTGGATT |
| AURKA | GCAGATTTTGGGTGGTCAGT | TCCGACCTTCAATCATTTCA |
| CDC20 | CTGTCTGAGTGCCGTGGAT | TCCTTGTAATGGGGAGACCA |
| DLG5 | ACCAGAAGGAGATCGGTGAC | ATCTCGGATGACCCGTTGT |
| IRF6 | TTTGTCTGGAAACATTCCTTAGC | CCCCAAAGCATAAGTAGATCTCAA |
| MINK1 | AGAAGCGGGGTGAGAAAGA | CGTTCATGATGGAGCTTGG |
| ΔNp63 | GAAAACAATGCCCAGACTCAA | TGCGCGTGGTCTGTGTTA |
| p53 | AGGCCTTGGAACTCAAGGAT | CCCTTTTTGGACTTCAGGTG |
| p63 | CCCCACCTCTGAACAAAATGA | GGGTTGATAAGCTGGCTCACA |
| SYNPO | AGGGAGGACCTAGCAGACG | GTCAGCTGGGCTGCAATC |
| TOP2A | TCTGGTCCTGAAGATGATGCT | TTAGTTAACCATTCCTTTCGATCA |
| TAZ | GTATCCCAGCCAAATCTCGT | TTCTGCTGGCTCAGGGTACT |
| YAP | GACATCTTCTGGTCAGAGATACTTCTT | GGGGCTGTGACGTTCATC |

**guide RNAs**

| Region | fw primer | bw primer |
|---|---|---|
| cdc20_Enhancer1_sgRNA_a | CACCGGATAGCAAACTGATTTCTGG | AAACCCAGAAATCAGTTTGCTATCC |
| cdc20_Enhancer1_sgRNA_b | CACCGGGCTGGTGCACGGATCTGCA | AAACTGCAGATCCGTGCACCAGCCC |
| cdc20_Enhancer2_sgRNA_c | CACCGTTACCCACATTATCAATGGT | AAACACCATTGATAATGTGGGTAAC |
| cdc20_Enhancer2_sgRNA_d | CACCGGGACGTGTAAGGGAGCTCTG | AAACCAGAGCTCCCTTACACGTCCC |
| cdc20_Enhancer2_sgRNA_e | CACCGAGTGAGTAATCCATGGCATG | AAACCATGCCATGGATTACTCACTC |
| sgRNA_control | CACCGAATCTCGCTTATATAACGAG | AAACCTCGTTATATAAGCGAGATTC |

**siRNAs**

| Gene | Name/ sequence | Reference |
|---|---|---|
| ctrl | Cat#4390843 | Thermo Fisher Scientific |
| YAP | UCUCUGACCAGAAGAUGUC | Azzolin Cell 2014; 158:157–70. |
| TAZ | ACGUUGACUUAGGAACUUU | Azzolin Cell 2014; 158:157–70. |

**Supplemental Table 2: Antibodies used in the study**

| | | |
|---|---|---|
| Mouse monoclonal anti-β-Actin | Santa Cruz Biotechnology | Cat# sc-47778; RRID: AB_626632 |
| Rabbit polyclonal anti-LIN9 | Bethyl | A300-BL2981 |
| Anti-mouse HRP conjugated | GE Healthcare | Cat# NXA931; RRID: AB_772209 |
| HRP Protein A | BD Biosciences | Cat# 610438; RRID: N/A |
| IgG from rabbit serum | Sigma | Cat# I5006; RRID: AB_1163659 |
| Mouse monoclonal anti CDK7 | Cell Signaling | Cat#2916; RRID:AB_2077142 |
| Mouse monoclonal anti Pol II 8WG16 | Santa Cruz Biotechnology | Cat#:sc-56767; RRID:AB_785522 |
| Mouse monoclonal anti Pol II CTD4H8 (Ser5) | Santa Cruz Biotechnology | Cat# sc-47701; RRID:AB_677353 |
| Mouse monoclonal anti-CDC20 | Santa Cruz Biotechnology | Cat# sc-13162; RRID: AB_628089 |
| Mouse monoclonal anti-p53 (DO-1) | Santa Cruz Biotechnology | Cat# sc-126; RRID:AB_628082 |
| Mouse monoclonal anti-p63 (D2K8X) | Cell Signaling | Cat#: 13109; RRID:AB_2637091 |
| Mouse monoclonal anti-YAP | Santa Cruz Biotechnology | Cat# sc-101199;RRID: AB_1131430 |
| Rabbit polyclonal anti-YAP | Novus Biologicals | NB110-58358 ; RRID:AB_922796 |
| Rabbit polyclonal anti acetyl Histone H4 (K5,8,12,16) | Merck | Cat#06-598 RRID:AB_310550 |
| Rabbit polyclonal anti CDK7 | Bethyl | Cat# A300-405A; RRID:AB_2275973 |
| Rabbit polyclonal anti Histone H3 (K27 acetyl) | Merck | Cat# 07-360; RRID: AB_310550 |
| Rabbit polyclonal anti Phospho-Chk1 (Ser345) | Cell Signaling | Cat#2341 ;RRID:AB_330023 |
| Rabbit polyclonal anti RNA Pol II- phospho ser2 | Abcam | Cat#ab5095; RRID:AB_304749 |
| Rabbit polyclonal anti RNA Pol II- phospho ser5 | Abcam | Cat# ab5131; RRID:AB_449369 |
